# Bovine breed-specific augmented reference graphs facilitate accurate sequence read mapping and unbiased variant discovery

**DOI:** 10.1101/2019.12.20.882423

**Authors:** Danang Crysnanto, Hubert Pausch

## Abstract

**Background:** The current bovine genomic reference sequence was assembled from the DNA of a Hereford cow. The resulting linear assembly lacks diversity because it does not contain allelic variation. Lack of diversity is a drawback of linear references that causes reference allele bias. High nucleotide diversity and the separation of individuals by hundreds of breeds make cattle ideally suited to investigate the optimal composition of variation-aware references.

**Results:** We augment the bovine linear reference sequence (ARS-UCD1.2) with variants filtered for allele frequency in dairy (Brown Swiss, Holstein) and dual-purpose (Fleckvieh, Original Braunvieh) cattle breeds to construct either breed-specific or pan-genome reference graphs using the *vg toolkit*. We find that read mapping is more accurate to variation-aware than linear references if pre-selected variants are used to construct the genome graphs. Graphs that contain random variants do not improve read mapping over the linear reference sequence. Breed-specific augmented and pan-genome graphs enable almost similar mapping accuracy improvements over the linear reference. We construct a whole-genome graph that contains the Hereford-based reference sequence and 14 million alleles that have alternate allele frequency greater than 0.03 in the Brown Swiss cattle breed. We show that our novel variation-aware reference facilitates accurate read mapping and unbiased sequence variant genotyping for SNPs and Indels.

**Conclusions:** We developed the first variation-aware reference graph for an agricultural animal: https://doi.org/10.5281/zenodo.3759712. Our novel reference structure improves sequence read mapping and variant genotyping over the linear reference. Our work is a first step towards the transition from linear to variation-aware reference structures in species with high genetic diversity and many sub-populations.

## Introduction

A reference sequence is an assembly of digital nucleotides that are representative for a species’ genetic constitution. Discovery and genotyping of polymorphic sites from whole-genome sequencing data typically involve reference-guided alignment and genotyping steps that are carried out successively [1]. Variants are discovered at positions where aligned sequencing reads differ from corresponding reference nucleotides. Long read sequencing and sophisticated genome assembly methods enabled spectacular improvements in the quality of linear reference sequences particularly for species with gigabase-sized genomes [2]. Recently generated *de novo* assemblies exceed in quality and continuity all current reference sequences [3, 4]. However, modifications and amendments to existing linear reference sequences causes shifts in their coordinates that require large efforts from the genomics community to make data compatible with updated reference sequences [5].

Domestication and selection for beef and milk production under various environmental conditions have led to the formation of more than thousand breeds of cattle (*Bos taurus*) with distinct genetic characteristics and high allelic variation within and between breeds [6]. The 1000 Bull Genomes Project discovered almost 100 million sequence variants that are polymorphic in 2,700 cattle from worldwide cattle breeds [7, 8]. Nucleotide diversity is higher in cattle than human populations [7, 9]. Yet, all bovine DNA sequences are aligned to the linear consensus reference sequence of a highly-inbred Hereford cow to facilitate reference-guided variant discovery and genotyping [10, 11]. A genome-wide alignment of DNA fragments from a *B. taurus* individual differs from the Hereford-based reference sequence at between 7 and 8 million single nucleotide polymorphisms (SNPs) and small (<50 bp) insertions and deletions (Indels) [12, 13]. The number of differences is higher for DNA samples with greater divergence from the reference [14, 15].

The bovine linear reference sequence lacks allelic variation and nucleotides that might segregate at high frequency in animals from breeds other than Hereford. Lack of allelic diversity is an inherent drawback of linear reference sequences because it causes reference allele bias. DNA sequencing reads that contain only alleles that match corresponding reference nucleotides are more likely to align correctly than DNA fragments that also contain non-reference alleles [16, 17]. Reads originating from DNA fragments that are highly diverged from corresponding reference nucleotides will either obtain low alignment scores, or align at incorrect locations, or remain un-mapped [18]. Reference bias compromises analyses that are sensitive to accurately mapped reads and prevents the precise estimation of allele frequencies [16, 19–21].

Graph-based [17, 22] and personalized reference genomes [5, 23] mitigate reference allele bias. Existing linear reference coordinates can serve as backbones for variation-aware genome graphs. Nodes in the graph represent alleles at sites of variation and edges connect adjacent alleles. Once a variation-aware genome graph contains all alleles at known polymorphic sites, every haplotype can be represented as a walk through the graph [24]. However, an optimal balance between graph density and computational complexity is key to efficient whole-genome graph-based variant analysis because adding sites of variation to the graph incurs computational costs. Recently, Pritt et al. [18] developed the FORGe software tool to prioritize variants for graph genomes. Their results provide a framework to build genome graphs that enable read mapping accuracy improvements over linear references at tractable computational complexity. A genome graph-based sequence analysis workflow is implemented in the *variation graph toolkit* (*vg*, https://github.com/vgteam/vg) [22]. The *vg toolkit* enables the mapping of sequence reads to variation-aware graphs that incorporate linear reference coordinates as a backbone. It also facilitates to augment genome graphs with genetic variants that have more complex topology (e.g., duplications, inversions and translocations) [25]. Graph-based references have been investigated primarily in humans and species with small genome sizes [17]. High nucleotide diversity and the separation of individuals by breeds make cattle an ideally suited species to investigate the optimal composition of reference graphs for gigabase-sized genomes.

Here, we investigate sequence read mapping and variant genotyping accuracy using variation-aware reference structures in cattle. Using sequence variant genotypes of 288 cattle from four dairy and dual-purpose breeds, we construct breed-specific augmented and pan-genome reference graphs using the *vg toolkit* [22]. We prioritize sequence variants to be added to the graphs and assess accuracy of read mapping for variation-aware and linear references (Figure 1). We show that breed-specific augmented and pan-genome graphs allow for significant read mapping accuracy improvements over linear reference sequences. We also construct a bovine whole-genome reference graph and show that unbiased and accurate sequence variant genotyping is possible from this novel reference structure. Together, we hope that our study can serve as a first step towards the transition from linear to variation-aware references in species with high genetic diversity and many sub-populations.

**Figure 1.**
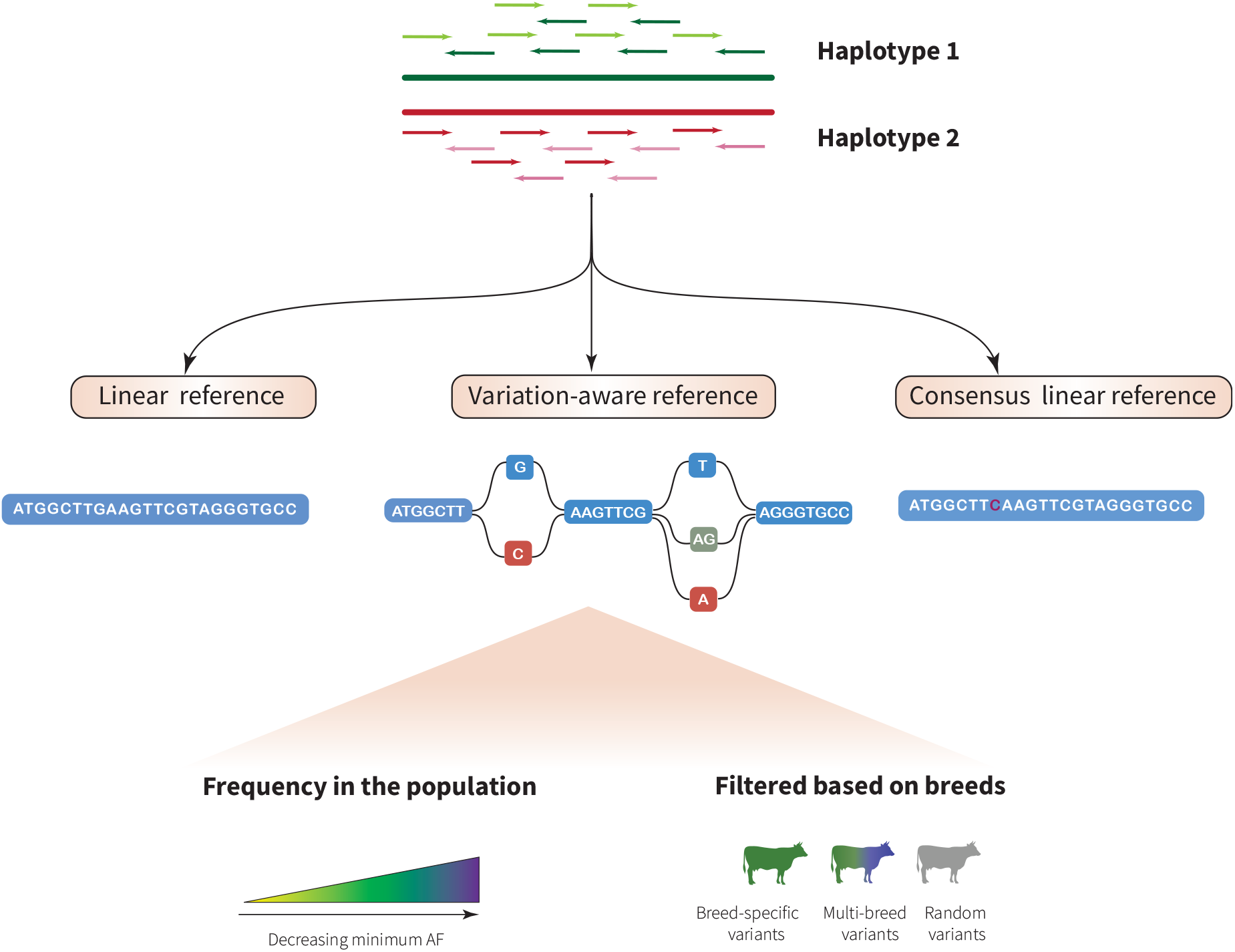
Schematic overview of the construction of breed-specific augmented genome graphs. We used the *vg toolkit* to augment the bovine linear reference sequence (ARS-UCD1.2) with alleles at SNPs and Indels that were discovered in 288 cattle from four breeds. Alleles that were added to the linear reference were prioritized based on their alternate allele frequency (AF). Reads simulated from true haplotypes were aligned to variation-aware, linear and consensus reference sequences to assess read mapping accuracy on cattle chromosome 25. Short-read sequencing data of Brown Swiss cattle were used to investigate sequence variant genotyping accuracy and reference allele bias using a bovine whole-genome graph as a novel reference.

## Results

### Construction of bovine breed-specific augmented genome graphs

Breed-specific augmented reference graphs were constructed for four genetically distinct dairy (Brown Swiss – BSW, Holstein – HOL) and dual-purpose (Fleckvieh – FV, Original Braunvieh – OBV) cattle breeds using the Hereford-based linear reference sequence (ARS-UCD1.2) of chromosome 25 as a backbone (Figure 2a). Average nucleotide diversity (*π*) estimated using 295,801 (HOL), 336,390 (FV), 347,402 (BSW) and 387,855 (OBV) biallelic variants of chromosome 25 ranged from 0.00177 (BSW) to 0.0019 (OBV) for the four breeds (Figure 2b). To determine the optimal composition of bovine variation-aware references, we augmented the linear reference of chromosome 25 with an increasing number of variants (SNPs and Indels) that were filtered for alternate allele frequency in 82 BSW, 49 FV, 49 HOL and 108 OBV cattle. In total, we constructed 20 variation-aware graphs for each breed that contained between 2,046 (variants had alternate allele frequency >0.9) and 293,804 (no alternate allele frequency threshold) alleles.

**Figure 2.**
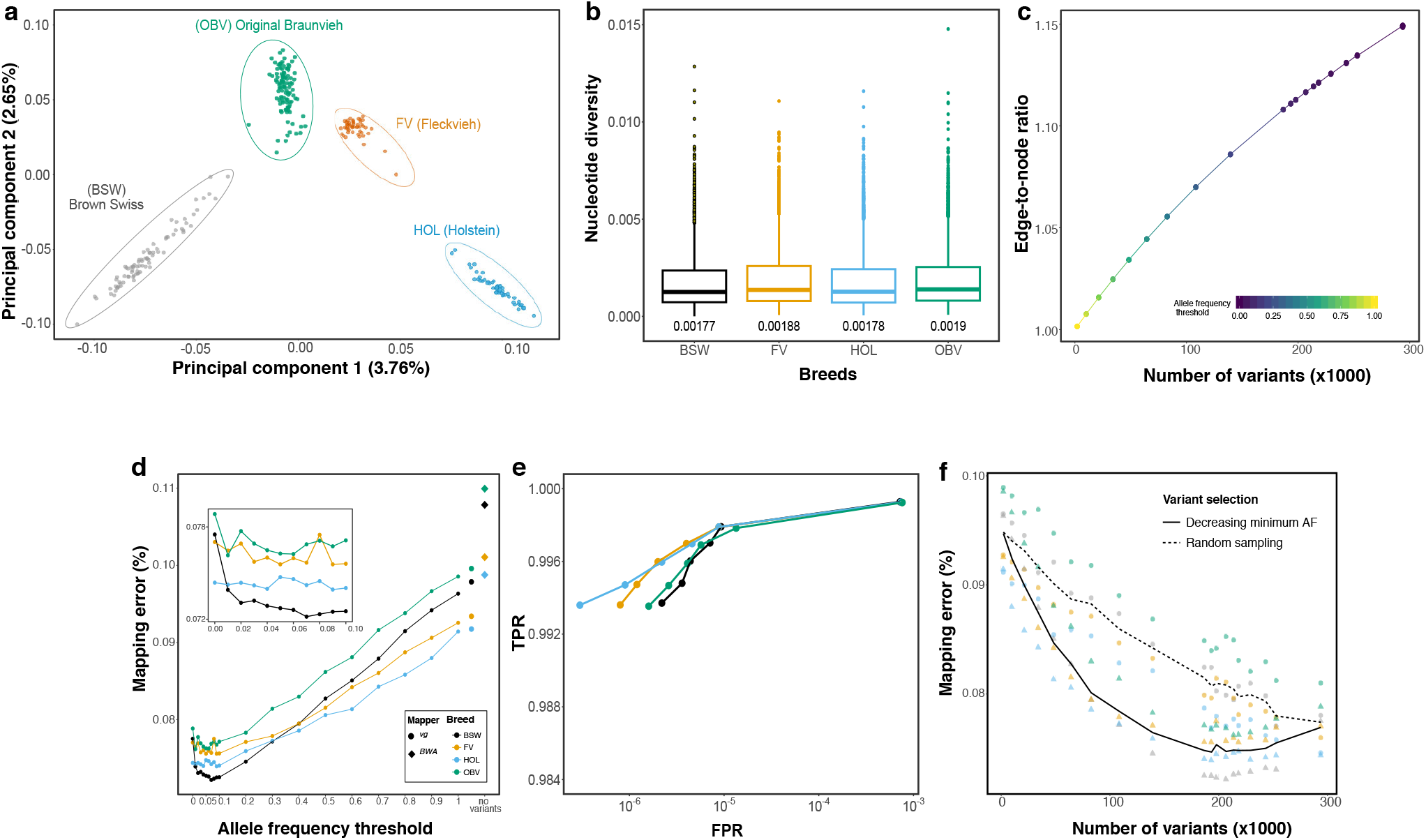
Accuracy of mapping simulated paired-end reads to genome graphs that contained variants filtered for allele frequency at chromosome 25. (a) The top principal components of a genomic relationship matrix constructed from whole-genome sequence variants reflect the genetic diversity of the four cattle breeds considered. (b) Nucleotide diversity of the four breeds calculated in non-overlapping 10 kb windows for variants of chromosome 25. The values below each boxplot indicate the nucleotide diversity for the four breeds averaged across all sliding-windows. (c) Edge-to-node ratio of graphs that contained between 2,046 and 293,804 variants filtered for allele frequency. (d) Proportion of incorrectly mapped reads for four breed-specific augmented genome graphs. Diamonds and large dots represent values from linear mapping using *BWA mem* and *vg*, respectively. The inset represents a larger resolution of the mapping accuracy for alternate allele frequency thresholds less than 0.1. (e) True positive (sensitivity) and false positive mapping rate (specificity) parameterized on mapping quality of the best performing graph from each breed. (f) Read mapping accuracy for breed-specific augmented graphs that contained variants that were either filtered for alternate allele frequency (triangles) or sampled randomly (circles) from all variants detected within a breed. The dashed and solid line represents the average proportion of mapping errors across four breeds using random sampling and variant prioritization, respectively. Colours indicate values obtained for different breeds. Results for single-end mapping are presented in Additional file 1: Figure S2.

The graph-based representation of bovine chromosome 25 (42,350,435 nucleotides) had 1,323,451 nodes and 1,323,450 edges. The number of nodes increased proportionally with the number of variants added to the reference. When we added a maximum number of 293,804 variants to the linear reference sequence of chromosome 25, the variation-aware graph contained 2.02 million nodes. The number of edges increased faster than the number of nodes, ranging from 1.32 (empty) to 2.33 (293,804 variants included) million. Consequently, the edge-to-node ratio increased when variants were added to the graph (Figure 2c). The number of paths through a graph grows rapidly with the number of variants being added to the graph. The index for the chromosome 25 reference graph contained 84.69 and 118.82 million k-mers (k=256) when 2046 and 293,804 variants, respectively, were added to the graphs (Additional file1: Figure S1).

### Variant prioritization based on allele frequency

We simulated 10 million paired-end reads (2×150 bp) corresponding to approximately 35-fold coverage of bovine chromosome 25 from haplotypes of BSW, FV, HOL and OBV cattle. Using either *BWA mem* or *vg*, we mapped the simulated reads to the respective breed-specific augmented reference graphs and the linear reference sequence. Variants that were only detected in animals used for read simulation were not added to the breed-specific augmented genome graphs. We observed fewer mapping errors using *vg* than *BWA mem* when simulated reads were aligned to a linear reference sequence. This finding was consistent for the four breeds investigated (Figure 2d). Variation-aware references that contained variants filtered for allele frequency in the respective breed reduced the mapping errors for all breeds. The proportion of reads with mapping errors decreased significantly with the number of variants added to the genome graph (Figure 2d, Pearson R=0.94, *P* < 10^−16^).

Read mapping accuracy increased almost linearly between alternate allele frequency threshold 1 and 0.1, i.e., until 186,680 variants with allele frequency greater than 0.1 were added to the graph (Pearson R=0.94, *P* < 10^−16^). Adding additional alleles that had alternate allele frequency between 0.1 and 0.01 to the graphs did not further improve read mapping accuracy over the scenario with an alternate allele frequency threshold of 0.1 (*P* = 0.13, Figure 2d inset). Read mapping accuracy declined (particularly in BSW) when the graphs contained rare alleles (alternate allele frequency<0.01) likely because such alleles are not observed in most animals of a population. Maximum read mapping accuracy was achieved at allele frequency thresholds between 0.2 and 0.01, when the graphs contained between 139,322 and 293,628 variants filtered for allele frequency. The number of erroneously mapped reads was clearly higher for graphs that contained randomly sampled than prioritized variants (Figure 2f). This finding corroborates that variant prioritization based on alternate allele frequency is important to achieve high mapping accuracy with graph-based reference structures.

We also applied the methods implemented in the FORGe software [18] to prioritize variants for the breed-specific augmented graphs (Additional file 2: Note S1). It turned out that genome graphs that were constructed with variants selected by the *Pop Cov* strategy, which relies solely on variant frequency information, enabled the highest mapping accuracy improvements over the linear reference. For example, we achieved the highest paired-end read mapping accuracy for the Brown Swiss reference graph (0.0722% erroneously mapped reads) using the *Pop Cov* method when we added 208,288 variants to the chromosome 25 reference (i.e., the top 60% of the ranked variants). The prioritized variants correspond to a minimum of alternate allele frequency of 0.06. Variant prioritization approaches that also take into account factors other than allele frequency e.g., the proximity of a variant to an already added variant in the graph or the repetitiveness of the resulting genome graph, did not lead to additional accuracy improvements.

Read mapping accuracy was highly correlated (Pearson R=0.94, *P* < 10^−16^) for single and paired-end reads (Additional file 1: Figure S2). However, the accuracy improvement of variation-aware over linear mapping was higher for single- than paired-end reads, possibly because distance and sequence information from paired reads facilitate linear read alignment.

Read mapping accuracy differed significantly among the four breeds analyzed (*P* = 10^−15^, linear model with allele frequency as covariate) although all breed-specific augmented graphs contained the same number of variants at each allele frequency threshold (Figure 2d). Linear mapping accuracy also differed among the breeds. We observed the highest error rate for reads aligned to the OBV-specific augmented reference graph. In 500 randomly sampled subsets of 35 sequenced cattle per breed, we discovered more sequence variants on chromosome 25 in OBV (N=305 ± 5K) than either FV (N=291 ± 3K), BSW (N=276 ± 6K) or HOL (N=259 ± 2K), reflecting that nucleotide diversity is higher in OBV than the other three breeds, which agrees with a recent study [26]. Across all alternate allele frequency thresholds considered, read mapping was more accurate for HOL than FV and OBV cattle, possibly because both genetic diversity and effective population size is less in HOL than the other breeds considered [27]. At allele frequency thresholds between 0.02 and 0.3, read mapping was more accurate for BSW than the other breeds. The proportion of variants with alternate allele frequency >0.02 was lower for BSW (84.1%) than the other breeds (86.3 – 89.2%). We detected more rare variants (allele frequency less than 0.05) in BSW and OBV than FV and HOL, likely reflecting differences in sample size (Additional file 1: Figure S3). An excess of singletons and rare variants in BSW and OBV cattle may have contributed to the decline in mapping accuracy at low alternate allele frequency thresholds (Figure 2d inset, Additional File 3: Table S2). Our findings indicate that differences in nucleotide diversity and allele frequency distributions across populations may affect read mapping accuracy to both linear and breed-specific augmented reference structures.

### Comparison between bovine and human genome graphs

We used publicly available whole-genome sequence variant data from phase 3 of the 1000 Genomes Project [28] to construct genome graphs for four genetically distinct human populations (Figure 3a, GBR (British, European), YRI (Yoruba Nigeria, African), STU (Sri Lankan Tamil, South Asia), and JPT (Japanese, East Asia)). The effective population size is more than 20-fold higher for the human than cattle populations (e.g. ~3100 for JPT and ~7500 for YRI [29] vs. ~80 for OBV and ~160 for FV [30, 31]). While the average number of sequence variants detected per sample was lower for the human (4,248,082) than cattle (6,973,036) populations, the proportion of singletons is higher in the human than cattle samples (23.00% in human vs. 14.01% in cattle) (Additional file 3: Table S1). The proportion of sequence variants that had minor allele frequency less than 0.05 was between 44.88% and 55.45% in the four human and between 23.65% and 38.70% in the four cattle populations (Additional file 1: Figure S4). Nucleotide diversity ranged from 0.00098 (JPT) to 0.00141 (YRI) (Figure 3b).

**Figure 3.**
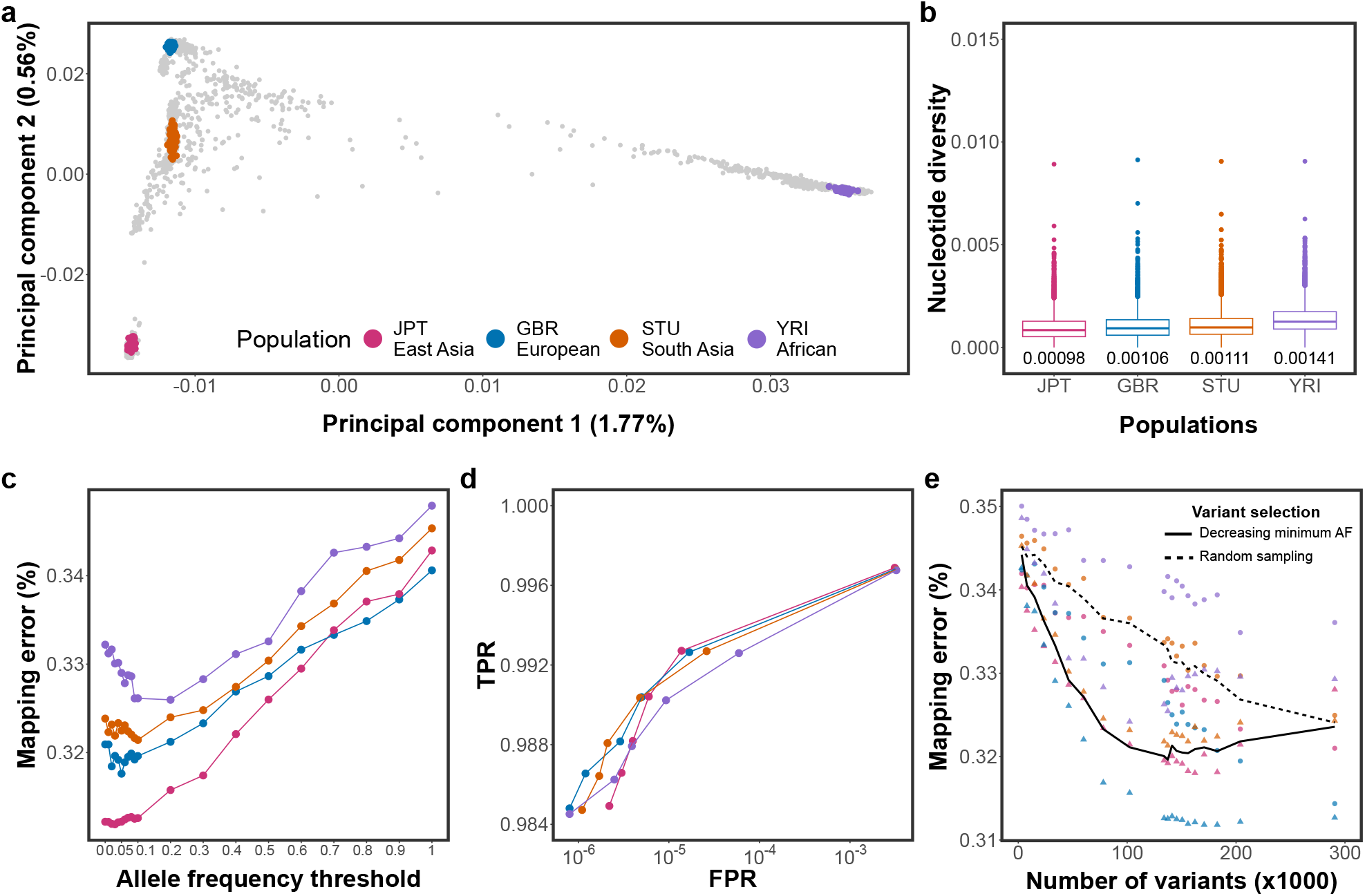
Accuracy of mapping simulated paired-end reads to human population-specific augmented genome graphs. (a) The top principal components of a genomic relationship matrix constructed from autosomal variants detected in 2,504 individuals that were included in phase 3 of the 1000 Genomes Project. The coloured points indicate 405 samples from the GBR (European), YRI (African), STU (South Asia), and JPT (East Asia) populations. (b) Nucleotide diversity of the four populations calculated in non-overlapping 10 kb windows for variants of chromosome 19. The values below each boxplot indicate the nucleotide diversity for the four populations averaged across all sliding-windows. (c) Proportion of incorrectly mapped reads for four population-specific augmented genome graphs. (d) True positive (sensitivity) and false positive mapping rate (specificity) parameterized on mapping quality of the best performing graph from each population. (f) Read mapping accuracy for population-specific augmented graphs that contained variants that were either filtered for alternate allele frequency (triangles) or sampled randomly (circles) from all variants detected within a population. The dashed and solid line represents the average proportion of mapping errors across four populations using variant prioritization and random sampling, respectively. Results for single-end mapping are presented in Additional file 1: Figure S6.

We considered the linear reference sequence of human chromosome 19 (g1k_v37 ref) as a backbone for the human genome graphs because its length (59,128,983 bp) and the number of variants detected per sample was similar to the values for bovine chromosome 25. Genetic diversity and allele frequency distributions were similar using either chromosome 19 or whole-genome variants indicating that the results obtained using chromosome 19 are representative for the human genome (Additional file1: Figure S4, S5, Additional file 3: Table S2). To construct population-specific augmented graphs, we used phased genotypes at 291,303, 306,304, 355,107, and 521,021 variants of chromosome 19 that were available for 104 JPT, 91 GBR, 102 STU and 108 YRI individuals. Once the variants that were only detected in individuals used for simulating reads were removed from the graphs, the population-specific augmented graphs for the GBR, YRI, STU and JPT populations contained between 3153 (alternate allele frequency >0.9) and 290,593 (no alternate allele frequency threshold) variants. We subsequently simulated 10 million reads from haplotypes of one individual per population and mapped the reads to the respective population-specific augmented genome graphs.

As observed for the bovine breed-specific augmented genome graphs, read mapping accuracy increased almost linearly between alternate allele frequency threshold 1 (no variants included) and 0.1 (133,891 variants added to the graph) (Figure 3c). Adding low-frequency variants (alternate allele frequency between 0.01 and 0.1) did not further improve the mapping accuracy. Mapping accuracy decreased for all graphs when we added very rare variants and singletons to the graphs. This pattern was most apparent for YRI which had the highest proportion of rare variants and nucleotide diversity among the four populations considered. Read mapping accuracy differed among the four populations analysed. We observed the lowest number of mis-mapped reads when reads simulated from a JPT individual were aligned to a JPT-specific augmented genome graph. The highest number of mis-mapped reads were observed when reads simulated from a YRI individual were aligned to a YRI-specific augmented genome graph. Mapping accuracy was higher for GBR than STU. These findings indicate that the mapping accuracy is negatively correlated with nucleotide diversity. Mapping accuracy improvements over the linear reference sequence were less when randomly sampled variants were added to the graphs (Figure 3e).

While the overall pattern of the mapping accuracy improvements over the linear reference was similar for human and bovine genome graphs across all allele frequency thresholds considered, the proportion of mis-mapped paired-end reads was approximately four-fold higher in the human than bovine alignments (two-fold for single-end reads; Additional file1: Figure S6). This finding was also apparent when the population-specific augmented graphs were parameterized on mapping quality to obtain sensitivity and specificity (Figure 2e and Figure 3d).

### Mapping to breed-specific augmented genome graphs

Next, we compared read mapping accuracy between bovine breed-specific augmented and pan-genome graphs (i.e., graphs that contained variants filtered for allele frequency across multiple populations) using reads simulated from phased variants of bovine chromosome 25. We constructed four breed-specific augmented genome graphs that contained variants that had alternate allele frequency >0.03 in either the BSW, FV, HOL or OBV breeds. HOL had the lowest number of variants (N=243,145) with alternate allele frequency >0.03, reflecting that sample size was lower in HOL than the other breeds. To ensure that the density of information was comparable across all breed-specific augmented graphs, we randomly sampled 243,145 variants with alternate allele frequency >0.03 from the BSW, FV and OBV populations and added them to the respective graphs. The pan-genome graph contained variants that had alternate allele frequency >0.03 in 288 individuals from the four populations. The random graph contained 243,145 randomly sampled variants for which haplotype phase and the allele frequency in the BSW, FV, HOL or OBV breeds was unknown (see Material and Methods section). To investigate read mapping accuracy, we simulated 10 million sequencing reads (150 bp) from BSW haplotypes and mapped them to the variation-aware and linear reference sequences. Variants that were only detected in the BSW animal used for simulating reads were excluded from the graphs. However, in order to determine an upper bound for graph-based read mapping accuracy, we also constructed a “personalized” genome graph, i.e., a graph that contains only haplotypes of the animal used for simulating the reads. We repeated the selection of variants, construction of variation-aware graphs and subsequent read mapping ten times.

The average length, number of nodes, number of edges and edge-to-node ratio of the variation-aware graphs was 42.60 Mb, 1,907,248, 2,155,799 and 1.13, respectively. Most variants of the random graph (87.81%) were not detected at alternate allele frequency >0.03 in BSW, FV, OBV and HOL indicating that they were either very rare or did not segregate in the four breeds considered in our study. Of 243,145 variants, an intersection of 48.13% had alternate allele frequency greater than 0.03 in the four breeds considered (Figure 4a). The average number of variants that were specific to the breed-specific augmented graphs ranged from 8,010 in BSW to 20,392 in FV.

**Figure 4.**
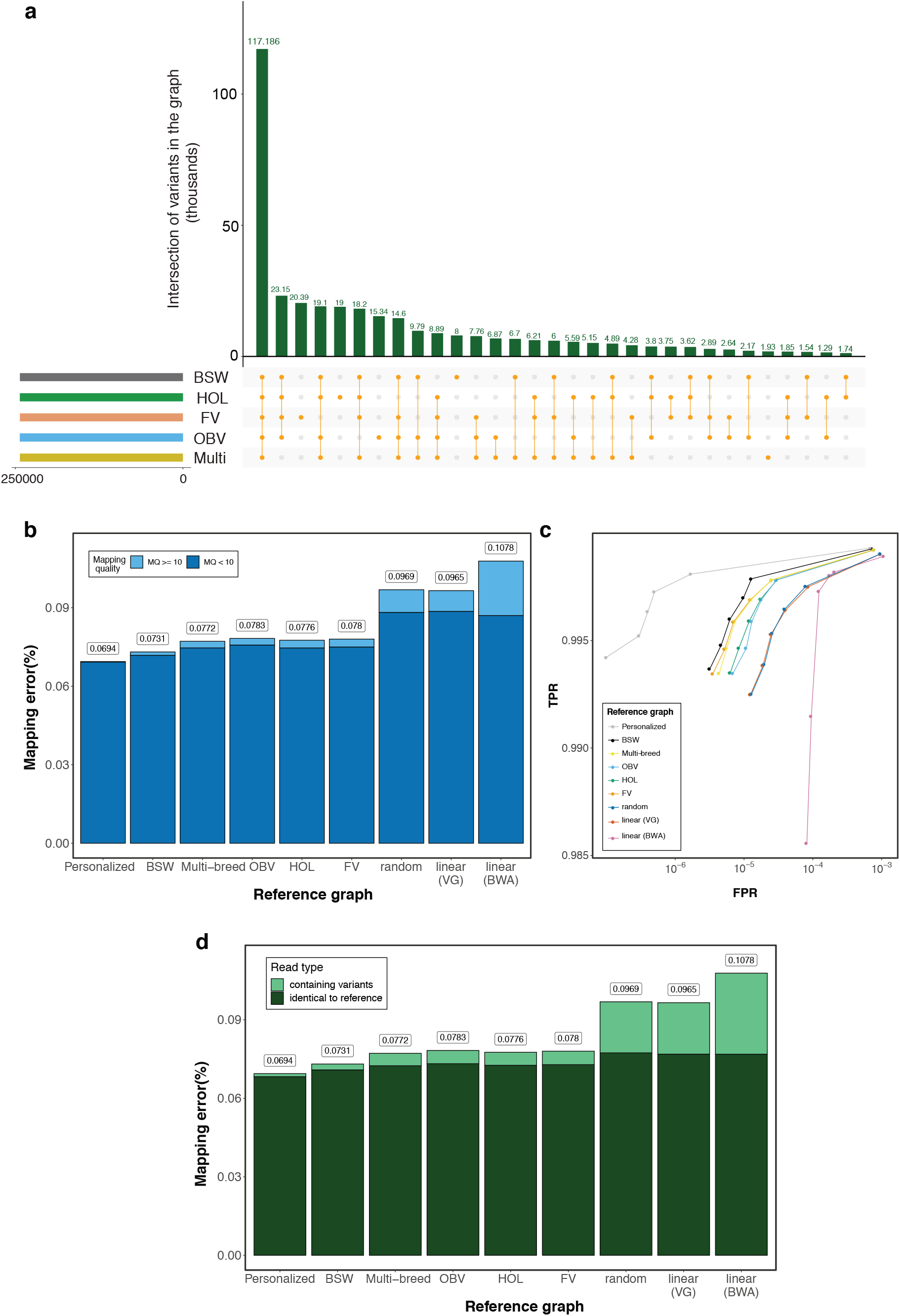
The accuracy of mapping simulated BSW paired-end reads to variation-aware and linear reference structures. (a) We added 243,145 chromosome 25 variants to the Hereford-based reference sequence that were filtered for alternate allele frequency >0.03 in either the BSW, FV, HOL or OBV populations. The pan-genome graph (Multi) contained 243,145 variants that had alternate allele frequency threshold >0.03 across 288 cattle from the four breeds considered. The bars indicate the overlap of variants (averaged across ten replications) that were added to different graphs (b) Proportion of simulated BSW reads that mapped erroneously against personalized graphs, breed-specific augmented graphs, pan-genome graphs (Multi-breed), random graphs or linear reference sequences. We used *vg* and *BWA mem* for linear mapping. Dark and light blue colours represent the proportion of incorrectly mapped reads that had phred-scaled mapping quality (MQ)<10 and MQ>10, respectively. (c) True positive (sensitivity) and false positive mapping rate (specificity) parameterized on mapping quality. (d) Proportion of BSW reads that mapped incorrectly against breed-specific augmented graphs, pan-genome graphs (Multi-breed), random graphs or linear reference sequences. Dark and light green colours represent the proportion of incorrectly mapped reads that matched corresponding reference nucleotides and contained non-reference alleles, respectively. Results for single-end mapping are presented in Additional file 1: Figure S7.

Personalized genome graphs, i.e., graphs that are tailored to a specific individual, enable the largest read mapping accuracy improvements over linear references. The proportion of mis-mapped reads was 0.0694% when a personalized BSW graph was used as a reference. Apart from the personalized graph, the highest mapping accuracy, sensitivity and specificity was achieved when the simulated BSW reads were aligned to a BSW-specific augmented graph (Figure 4bcd). The proportion of erroneously mapped paired-end reads was 0.073% for the BSW-specific augmented graph. Sensitivity and specificity were slightly lower and the number of reads with mapping errors was slightly higher when the same reads were aligned to a pan-genome graph. The read mapping accuracy differed only slightly between the breed-specific augmented and pan-genome graph because the overlap of variants that were included in both variation-aware references was high (Additional file 1: Figure S8). The number of mapping errors was higher (adjusted P < 10^−16^, pairwise t test, Additional file 1: Figure S9) when BSW reads were aligned to genome graphs that contained variants filtered for allele frequency in either the FV, HOL or OBV populations.

We also simulated reads from haplotypes of FV, HOL and OBV cattle. Similar to our findings using reads simulated from BSW cattle, mapping was more accurate to breed-specific than either pan-genome graphs or graphs that were augmented with variants filtered for allele frequency in other breeds (Additional File 1: Figure S10).

Mapping reads to a linear reference sequence using *BWA mem* with default parameter settings was the least sensitive and least specific mapping approach tested. Linear mapping using *vg* was also less accurate than variation-aware mapping. This finding indicates that accuracy improvements of variation-aware over linear mapping are attributable to differences in the reference structure rather than mapping algorithms. All graphs that contained pre-selected variants that had alternate allele frequency greater than 0.03 enabled significantly (P=10^−16^, two-sided t test) higher mapping accuracy than linear references. This was also true when reads were mapped to graphs that contained variants that were filtered for allele frequency in a different breed, likely because many common variants segregated across the four breeds considered (Figure 4a).

Recently, Grytten et al. [32], showed that an adjusted linear alignment approach that relies on a combination of *BWA mem* and *Minimap2* [33] may improve linear mapping accuracy because the default setting of *BWA mem* might miss sub-optimal alignments and overestimate mapping quality for multi-mapping reads [32, 34]. We found that this approach enables to reduce the proportion of mis-mapped from 0.1078 to 0.0983 in cattle (Additional file 2: Note S2). Improved mapping accuracy from the combination of *BWA mem* and *Minimap2* primarily results from less incorrectly mapped reads that had mapping quality >10, indicating a better mapping quality assignment. The mapping accuracy from the adjusted linear alignment approach is similar to the linear mapping accuracy obtained using *vg* but considerably lower than using breed-specific augmented graphs (Additional file 2: Note S2). The number of paired-end reads with mapping errors is 26% higher using the adjusted linear alignment approach than breed-specific augmented reference graphs.

Reference graphs that contained random variants, i.e., variants that were neither phased, nor filtered for allele frequency in the breeds of interest, did not improve mapping accuracy, sensitivity and specificity over linear references (adjusted P = 0.74 and 0.35 for single- and paired-end, pairwise t-test, Additional file 1: Figure S9).

Compared to linear mapping using *BWA mem* with default parameter settings, the number of mapping errors decreased by 39 and 31% for single- and paired-end reads, respectively, using a breed-specific augmented reference graph. Extrapolated to whole-genome sequencing data required for a 35-fold genome coverage, the use of breed-specific augmented reference graphs could reduce the number of incorrectly mapped single- and paired-end reads by 1,3000,000 and 220,000, respectively.

Using the BSW-specific augmented graph as a reference, only 1.76% of the incorrectly mapped reads had mapping quality (MQ) greater than 10. The MQ of the vast majority (98.24%) of incorrectly mapped reads was less than 10, i.e., they would not qualify for sequence variant discovery and genotyping using *GATK* with default parameter settings. The proportion of incorrectly mapped reads with MQ>10 was twice as high using either the pan-genome or an across-breed augmented reference graph (3.21-3.85%). The proportion of incorrectly mapped reads with MQ>10 was higher using either the random graph (8.92%) or linear reference sequence (*vg*: 8.19%, *BWA mem*: 19.3%).

Of 10 million simulated reads, 19.16% contained at least one nucleotide that differed from corresponding Hereford-based reference alleles. Using *BWA mem*, 47.44% and 28.72% of the erroneously mapped single- (SE) and paired-end (PE) reads, respectively, contained alleles that differed from corresponding reference nucleotides indicating that incorrectly mapped reads were enriched for reads that contained non-reference alleles (Figure 4d, Additional file 1: Figure S7, Additional file 1: Figure S11). The proportion of erroneously mapped reads that contained non-reference alleles was similar for reads that were aligned to either random (47.62% and 20.13%) or empty graphs (48.20% and 20.35%) using *vg*. However, the proportion of incorrectly mapped reads that contained non-reference alleles was clearly lower for the breed-specific augmented (SE: 1.37%, PE: 3.08%) and pan-genome graphs (SE: 2.12%, PE: 6.14%). The proportion of incorrectly mapped reads that matched corresponding reference nucleotides was almost identical across all mapping scenarios tested (Figure 4d, Additional file 1: Figure S7, Additional file 1: Figure S11).

Using data from the Ensembl bovine gene annotation (version 99) and RepeatMasker, we determined if the simulated reads originate from either genic regions, interspersed duplications or low-complexity and simple repetitive regions (Additional file 1: Figure S12). Regardless of the reference structure used, the mapping accuracy was low for reads originating from repetitive regions. Mapping accuracy was higher for reads originating from either genic or exonic regions. Graph-based references enabled more accurate mapping of reads originating from either genic regions or interspersed duplications (including SINEs, LINEs, LTR, and transposable elements) than linear reference sequences. However, graph-based references did not improve the mapping accuracy over linear references for reads that originate from low-complexity or simple repetitive regions.

We further augmented the BSW-specific genome graph with 157 insertion and deletion polymorphisms of bovine chromosome 25 that were detected from short paired-end reads (2×150 bp) of 82 BSW animals using *Delly*. Adding these variants to the graph either alone or in addition to 243,145 variants that were detected using *GATK* did not improve the mapping accuracy over the corresponding scenarios that did not include these variants (Additional file 2: Note S3).

### Linear mapping accuracy using a consensus reference sequence

Previous studies reported that linear mapping may be more accurate using population-specific than universal linear reference sequences [5, 35, 36]. In order to construct bovine linear consensus reference sequences, we replaced the alleles of the chromosome 25 ARS-UCD1.2 reference sequence with corresponding major alleles at 67,142 and 73,011 variants that were detected in 82 BSW and 288 cattle from four breeds, respectively. Subsequently, we aligned 10 million simulated BSW reads to the linear adjusted sequences using either *vg* or *BWA mem*. Read mapping was more accurate to the consensus than original linear reference sequence (Figure 5, Additional file 1: Figure S13). The accuracy of mapping was higher when reference nucleotides were replaced by corresponding major alleles that were detected in the target than multi-breed population. However, the mapping of reads was less accurate, sensitive and specific using either of the consensus linear reference sequences than the breed-specific augmented graphs (Figure 5b).

**Figure 5.**
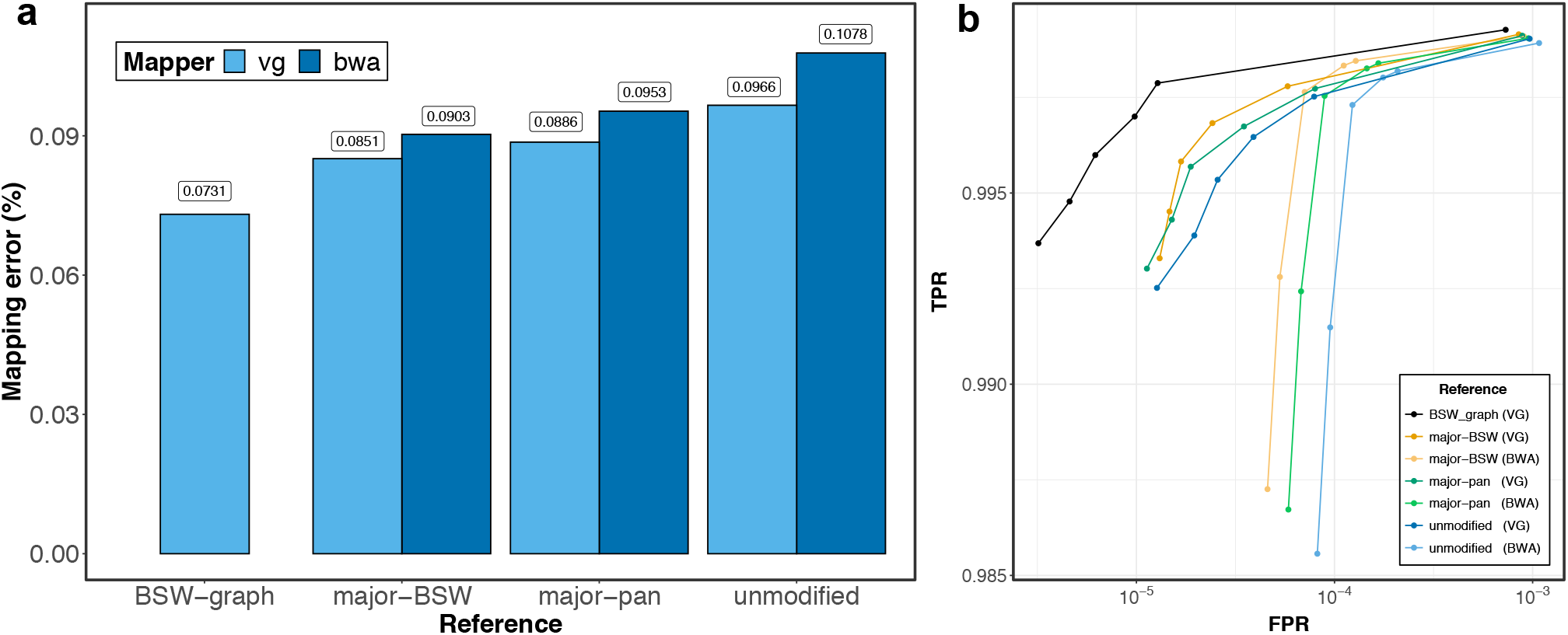
Paired-end read mapping accuracy using breed-specific augmented genome graphs and consensus linear reference sequences. (a) Dark and light blue represent the proportion of reads that mapped incorrectly using *BWA mem* and *vg*, respectively, to the BSW-specific augmented reference graph (BSW-graph), the BSW-specific (major-BSW) and the multi-breed linear consensus sequence (major-pan) and the bovine linear reference sequence (unmodified). (b) True positive (sensitivity) and false positive mapping rate (specificity) parameterized based on the mapping quality. The results of an analysis where reference nucleotides were only replaced at SNPs is available in Additional file 1: Figure S13.

### Read mapping and variant genotyping using whole genome graphs

In order to develop a breed-specific augmented reference structure for whole-genome applications, we constructed a BSW-specific augmented whole-genome variation-aware reference graph using 14,163,824 autosomal biallelic variants (12,765,895 SNPs and 1,397,929 Indels) that had alternate allele frequency greater than 0.03 in 82 BSW cattle. The resulting graph contained 111,511,367 nodes and 126,058,052 edges (an edge-to-node ratio of 1.13) and 6.32 × 10^9^ 256-mer paths. We also constructed a linear (empty) whole-genome graph that did not contain allelic variation. Subsequently, we mapped paired-end (2×150 bp) sequencing reads of 10 BSW cattle that had been sequenced at between 6 and 40-fold coverage (Additional file 3 Table S4) to the variation-aware and linear reference sequence using either *vg map* or *BWA mem*. The 10 BSW cattle used for sequence read mapping were different to the 82 animals used for variant discovery, graph construction and haplotype indexing.

62.19, 51.35 and 49.16% of the reads aligned perfectly (i.e., reads that aligned with full length (no clipping) and without any mismatches or Indels) to the BSW-specific augmented graph, the empty graph, and the linear reference sequence, respectively (Figure 6a). We observed slightly less uniquely mapped reads using either the whole-genome (82.46%) or empty graph (82.18%) than the linear reference sequence (83.18%) indicating that variation-aware references can increase mapping ambiguity due to providing alternative paths for read alignment.

**Figure 6.**
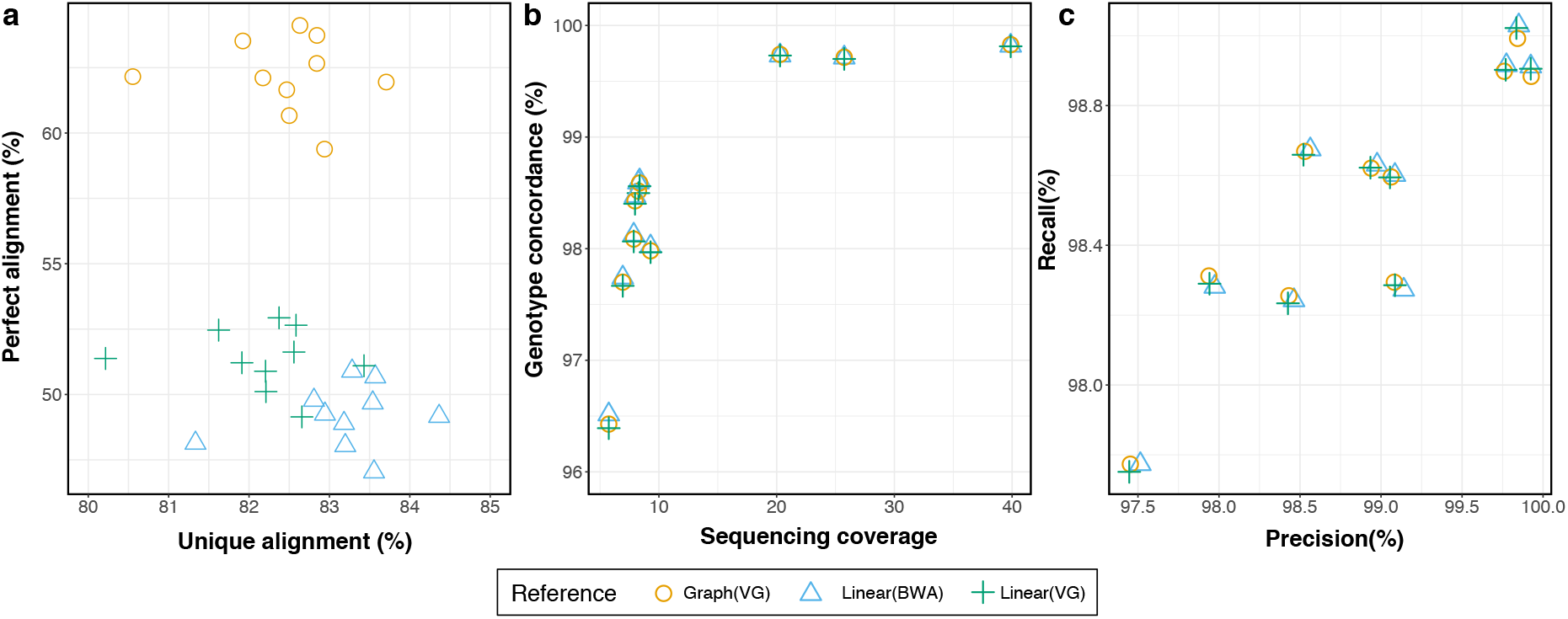
Sequence read mapping and variant genotyping using a breed-specific augmented whole-genome graph. (a) Proportion of sequencing reads that mapped perfectly and uniquely to the BSW-specific augmented (circle) and Hereford-based linear (triangle, cross) reference. (b) Concordance between sequence variant and corresponding microarray-derived genotypes as a function of sequencing depth. Sequence variant genotypes were obtained using the multi-sample variant calling approach implemented in *SAMtools*. (c) Corresponding precision-recall statistic. Each symbol represents one BSW animal.

We converted (surjected) the graph-based read alignments of 10 BSW cattle to corresponding linear reference coordinates and genotyped polymorphic sites using *SAMtools mpileup*. In order to assess genotyping accuracy, we compared the sequence variant genotypes with array-called genotypes at corresponding positions. Sequence variant genotyping accuracy was correlated with sequencing coverage (Figure 6b). Genotype concordance, non-reference sensitivity, non-reference discrepancy, and precision did not differ between the graph-based and linear alignments for both raw and hard-filtered genotypes (Figure 6bc, Additional file 3: Table S3). The average concordance, precision and recall from the graph-based alignments was 99.76, 99.84 and 98.93, respectively, for three samples (SAMEA6163185, SAMEA6163188, SAMEA6163187) with sequencing coverage greater than 20-fold. We observed similar values for genotypes called using either *GATK* or *Graphtyper* (Additional file 3: Table S3). In agreement with our previous findings [12], genotype concordance was slightly higher using *Graphtyper*, than either *SAMtools or GATK*.

### Variation-aware alignment mitigates reference allele bias

To investigate reference allele bias in genotypes called from linear and graph-based alignments, we aligned sequencing reads of a BSW animal that was sequenced at 40-fold coverage (SAMEA6163185) to either the BSW-specific augmented whole-genome graph or linear reference sequence (Additional file 3: Table S4). We called genotypes using either *SAMtools mpileup* or *GATK*. The genotypes were filtered stringently to obtain a high-confidence set of 2,507,955 heterozygous genotypes (2,217,069 SNPs and 290,886 Indels, see Material and Methods) for reference allele bias evaluation. The BSW-specific augmented whole-genome reference graph contained the alternate alleles at 2,194,422 heterozygous sites (87.49%).

Using *SAMtools* to genotype sequence variants from variation-aware and linear alignments, the support for reference and alternate alleles was almost equal at heterozygous SNPs (Figure 7a), indicating that SNPs are not notably affected by reference allele bias regardless of the reference structure. Alternate allele support decreased with variant length for the linear alignments. As expected, bias towards the reference allele was more pronounced at insertion than deletion polymorphisms. For instance, for 456 insertions that were longer than 30 bp, only 26% of the mapped reads supported the alternate alleles. The allelic ratio of Indel genotypes was closer to 0.5 using graph-based than linear alignments indicating that variation-aware alignment mitigates reference allele bias. However, slight bias towards the reference allele was evident at insertions with length >12 bp, particularly if the alternate alleles were not included in the graph (Figure 7a). Inspection of the read alignments using the *Sequence Tube Map* graph visualization tool [37] corroborated that the support for alternate alleles is better using graph-based than linear references (Additional file 1: Figure S14).

**Figure 7.**
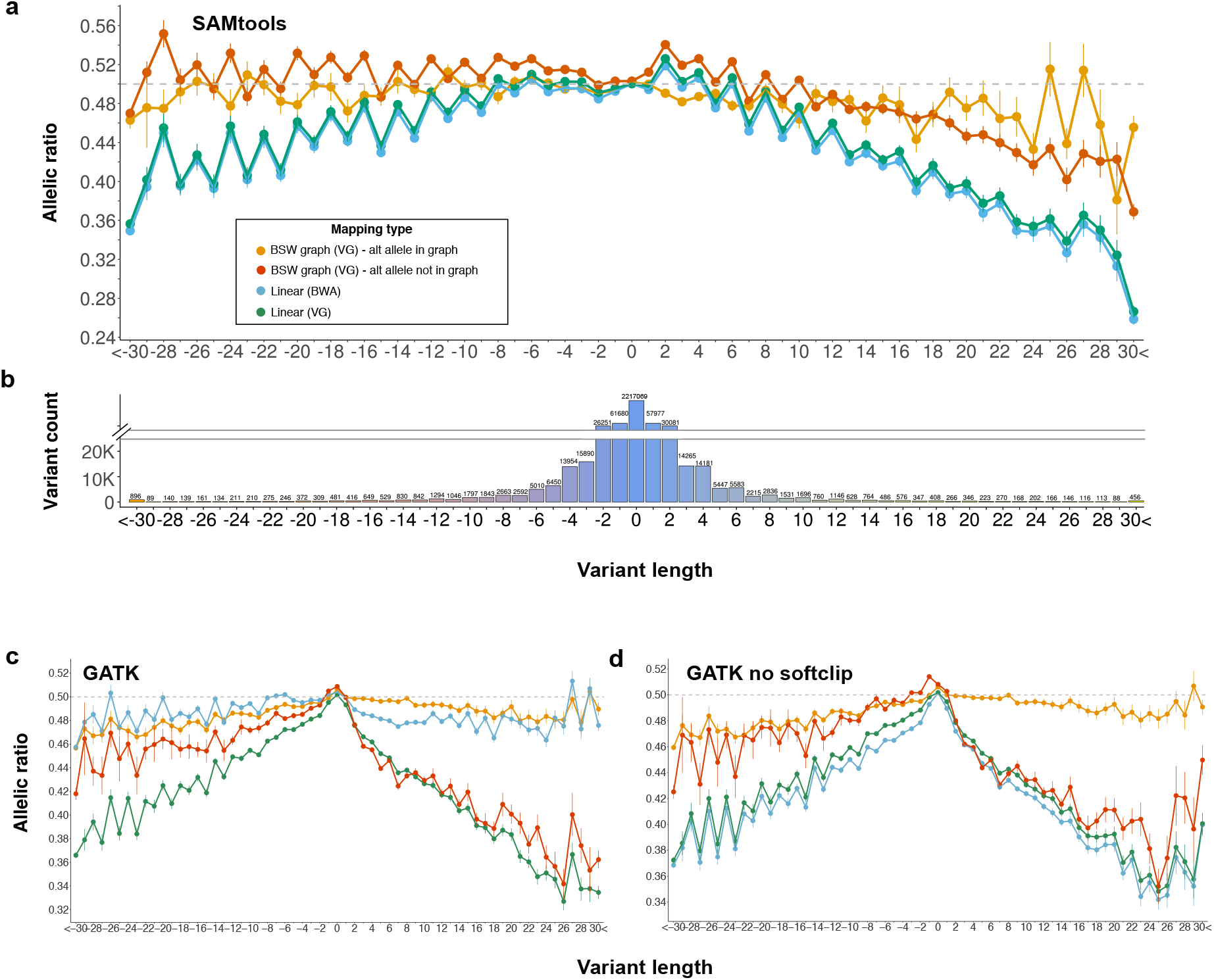
Reference allele bias from graph-based and linear alignments. Reference allele bias from graph-based and linear alignments using **(a)** *SAMtools*, **(c)** *GATK*, or **(d)** *GATK* without soft-clip for variant genotyping and either *BWA mem* or *vg* for alignment. Allelic ratio reflects the proportion of mapped reads supporting the alternate allele. The grey dashed line indicates equal support (0.5) for both alleles. Negative values, zero and positive values along the x axis represent deletions, SNPs, and insertions respectively. Each dot represents the mean (± s.e.m.) allelic ratio for a given variant length. **(b)** Number of variants with a given length. To improve the readability, the values above the breakpoint of the y-axis do not scale proportionately with the height of the bars

Both the number of reads mapped and the number of mapped reads supporting alternate alleles was higher at Indels using graph-based than linear alignments (Additional file 1: Figure S15). The difference in the number of mapped reads between graph-based and linear alignments increased with variant length. However, the number of mapped reads supporting the reference alleles did not differ between the graph-based and linear alignments. This finding indicates that reduced reference allele bias at Indel genotypes called from graph-based alignments is due to the improved mapping of reads that contain non-reference alleles.

We next investigated if these conclusions also hold for genotypes called by *GATK*. While *SAMtools mpileup* detects variants directly from the aligned reads [38], *GATK HaplotypeCaller* locally realigns the reads and calls variants from the refined alignments [39]. Using *GATK*, the allelic ratio was close to 0.5 for genotypes called from graph-based alignments across different lengths of variants that were included in the reference graph (Figure 7c). However, reference allele bias was evident at insertions that were not included in the reference graph. We also observed an almost equal number of reference and alternate alleles at variants genotyped from linear alignments using *GATK*. These findings confirm that the local realignment and haplotype-based genotyping approach of *GATK* might also mitigate reference alleles from linear alignments.

The percentage of soft-clipped reads increased with Indel length in the linear alignments (Additional file 1: Figure S16). However, the graph-based alignments contained almost no soft-clipped reads across all Indel lengths. In order to investigate the impact of soft-clipping on variant genotyping, we repeated *GATK* variant discovery and genotyping for the graph-based and linear alignments after all soft-clipped reads were removed (Figure 7d). As expected, the allelic ratio of genotypes called from the graph-based alignments was not affected by the removal of (very few) soft-clipped reads. However, bias towards the reference allele became evident in genotypes called from linear alignments. This finding confirms that the local realignment of *GATK* rescues Indels that are initially soft-clipped, thus mitigating reference allele bias. This finding also implies that the original pileup information from graph-based alignments facilitates to confidently detect known Indels while avoiding local realignment as implemented in the *GATK HaplotypeCaller*.

## Discussion

To the best of our knowledge, our study is the first to investigate the utility of a variation-aware reference for a species with a gigabase-sized genome other than human. We constructed bovine breed-specific consensus sequences and variation-aware reference graphs using a Hereford-based linear reference sequence as backbone and variants that were filtered for allele frequency in four cattle breeds other than Hereford to investigate read mapping accuracy and variant genotyping from different reference structures.

Using sequencing reads simulated from haplotypes of BSW, FV, OBV and HOL cattle, our findings confirm that a breed-specific consensus sequence improves linear mapping [5, 35]. However, read mapping is less accurate using linear consensus than variation-aware references that contain pre-selected variants. Grytten et al. [32] reported that an adjusted parameter setting of *BWA mem* and subsequent application of *Minimap2* may further improve the linear mapping accuracy. However, the adjusted linear mapping approach still performs worse than graph-based mapping on reads that contain variants. The accuracy improvements of the adjusted linear mapping approach were small in our study, because the number of sequence variants detected per sample and thus the proportion of reads with variants is almost twice as high in cattle than humans (Additional file 3: Table S1).

Using a bovine variation-aware reference reduced the proportion of erroneously mapped reads by more than 30% compared to the most widely-used linear mapping approach. A similar improvement in mapping accuracy over the linear reference was achieved for a human variation-aware reference genome [18]. The graph-based alignments using the most accurate breed-specific augmented reference graph contained 0.073% erroneously mapped reads. Incorrectly mapped reads that had high mapping quality (MQ >10) were less frequent in the graph-based than linear alignments. Thus, a variation-aware reference may reduce the number of flawed genotypes arising from mapping errors that would remain unnoticed due to high mapping quality. Similar to findings in human genome graphs [18, 25], bovine variation-aware references did not improve the mapping of short reads that originate from low-complexity regions.

Our findings demonstrate that variant prioritization is key to accurate variation-aware read mapping. Based on investigations in four genetically distinct cattle breeds and human populations, we make three important observations: first, variation-aware references that contain random variants for which the allele frequency and haplotype phase in the target populations is unknown, do not improve read mapping accuracy over linear references. Our previous study also showed that adding many random variants does barely affect sequence variant genotyping from reference graphs [12]. Adding random unphased variants increases the number of alternative alignment paths that are not necessarily biologically plausible haplotypes, thus increasing mapping ambiguity. Second, read mapping accuracy increases approximately linearly with the number of randomly sampled breed-specific variants being added to the genome graph. Similar findings in the four human population-specific augmented graphs confirm that this observation also holds for populations that are strongly enriched for rare alleles and singletons. Third, the highest mapping accuracy at tractable graph complexity can be achieved when variants filtered for allele frequency are added to the graph. Using variant prioritization approaches that are based on allele frequency, we observed the highest mapping accuracy at allele frequency thresholds between 0.01 and 0.10 in four cattle breeds and four human populations. In order to reduce the computational complexity of variation-aware read mapping, previous studies used arbitrarily chosen allele frequency thresholds to prioritize variants to be included in the graphs (e.g., 1% [22, 40], 5% [41], 10% [42]). Using fine-grained allele frequency inclusion thresholds, we find that the read mapping accuracy does not notably differ between the 0.01 and 0.1% thresholds in most populations. Yet, mapping accuracy declined rapidly for the YRI-specific augmented graph when variants with frequency less than 10% were added indicating that the optimal inclusion threshold may vary across populations. Variant prioritization approaches that also take into account factors other than allele frequency [18] did not lead to further accuracy improvements in our study. Considering that most cattle breeds have an effective population size between 50 and 200 [43, 44], the vast majority of variants with allele frequency greater than 0.1 can be detected from a few sequenced key ancestor animals [13]. As a matter of fact, key ancestor animals have been sequenced for many cattle breeds [7, 45]. Thus, the construction of variation-aware reference structures that are informative for many cattle breeds is readily possible using e.g., the sequence variant catalogue of the 1000 Bull Genomes Project [7, 8].

A pan-genome graph that contained variants filtered for allele frequency across the four cattle breeds enabled almost similar accuracy improvements over the linear reference than breed-specific augmented graphs (Figure 4b). Although the principal components analysis confirmed that the breeds considered in our study are genetically distinct populations, they share many common alleles. Moreover, compared to human populations, the proportion of rare alleles and singletons is low in cattle. The bovine pan-genome graph constructed in our study contained between 75.28 and 80.82% of the variants that were also added to the breed-specific augmented graphs. Instead of building many breed-specific graphs, the construction of a universal pan-genome graph is likely possible without notably compromising the accuracy of read mapping. This conclusion may hold for many species with genetically distinct sub-populations that share common alleles. Compared to the linear reference, the mapping accuracy was also significantly higher when reads from one breed were mapped to a genome graph that contained variants filtered for allele frequency in another somewhat related breed. Thus, the BSW-specific augmented whole-genome graph constructed in our study will likely improve read mapping accuracy over the linear reference and mitigate reference allele bias also for breeds other than BSW, FV, HOL and OBV. Our BSW-specific augmented whole-genome graph is available at https://doi.org/10.5281/zenodo.3759712 [46]. In order to facilitate the construction of variation-aware reference structures, the entire workflow to establish whole-genome graphs is also available at https://github.com/danangcrysnanto/bovine-graphs-mapping.

The number of sequencing reads that aligned to the BSW-specific whole-genome graph with full identity increased considerably (+13%) over the linear reference sequence at the cost of a slightly reduced (−0.72%) number of unique alignments. A two-step graph alignment approach that exploits a refined search space might reduce the number of multiple mappings in dense variation-aware graphs [32]. Compared to a human whole-genome graph, the improvement in perfect mapping over the linear reference was slightly larger in our bovine whole genome graph (9.2%) [22]. However, the proportion of reads with perfect alignments (62.19%) was lower in our BSW-specific whole-genome graph, likely because it contained only sequences that were assembled to the 29 autosomes. The graph did not contain 269.77 Mb of the sex chromosomes, mitochondrial DNA, and 2180 unplaced contigs. A more sophisticated assembly of the bovine genome with increased continuity particularly at the sex chromosomes [4, 47] might serve as a backbone for an improved variation-aware genome graph.

In order to detect SNPs and Indels from the variation-aware reference graph using widely-used sequence variant genotyping methods, we had to make the graph-based alignments compatible with linear coordinates. Thus, our assessment of sequence variant genotyping from the bovine whole-genome graph is based on surjected graph-based alignments. It is possible that converting graph-based to linear alignments compromises variant discovery. However, the accuracy and sensitivity of genotyping did not differ between graph-based and linear alignments indicating that our whole-genome graph facilitates accurate sequence variant (SNPs and small Indels) genotyping. It is worth noting that our analysis considered only SNPs that are located in well-accessible regions of the genome, thus possibly overestimating genotyping accuracy [48, 49]. A benchmark dataset that enables unbiased evaluation of sequence variant genotyping [50] is not available for the four cattle breeds considered in our study. Because approximately 90% of the considered SNPs were already included in the BSW-specific whole-genome graph, they can be detected and genotyped easily from graph-based alignments [17]. These variants can also be detected and genotyped accurately from linear alignments [12, 51].

As expected, bias towards the reference allele was less in graph-based than linear alignments particularly at variants that were included in the graph. Unbiased genotyping of heterozygous variants from graph-based alignments is possible because reads supporting alternate alleles align better to variation-aware than linear references. Thus, our bovine whole-genome graph offers an appealing novel reference for investigations that either rely on low-coverage sequencing or are sensitive to unbiased allele frequencies [16, 19, 52]. Because a benchmark dataset for an unbiased evaluation of sequence variant genotyping performance [50] is not available in cattle, our assessment was restricted to heterozygous variants that were identified from both linear and graph-based alignments. This set of variants is possibly enriched for variants that can be called confidently from linear alignments, thus underestimating the graph-based genotyping performance (e.g., [22]).

Our study has three limitations. First, variants used to construct the breed-specific augmented genome graphs might be biased because they were detected from linear alignments of short sequencing reads. Variant discovery from an independent variation-aware reference structure might allow for a more complete assessment of genetic variation [50]. Second, we used the Hereford-based linear reference sequence as backbone to construct breed-specific augmented reference sequences. However, the Hereford-based reference sequence might lack millions of basepairs that segregate in the four breeds considered in our study [53–56]. These nucleotides are likely missing in the breed-specific augmented reference graphs constructed in our study. Accurate and continuous genome assemblies from BSW, FV, HOL and OBV cattle are not available. All bovine genome assemblies that are available to date had been compiled from individuals that are distantly related to the breeds in our study [2, 4, 57]. Haplotype-resolved genome assemblies of cattle from different breeds will facilitate the construction of more informative genome graphs and make non-reference sequences and their sites of variation amenable to genetic investigations [2, 4]. Third, we did not investigate the impact of large sequence variation on sequence read mapping and variant genotyping performance because neither a high-quality benchmark set of large structural variants (cf. [58]) nor long-read sequencing data is available for the four cattle breeds considered. Adding insertion and deletion polymorphisms detected from short-read sequencing data did not lead to accuracy improvements in our study likely because structural variants detected from short reads are notoriously biased and incomplete [59]. Recent studies indicated that large structural variants can be identified accurately from genome graphs [25, 60–62]. Eventually, a bovine genome graph that unifies multiple breed-specific haplotype-resolved genome assemblies and their sites of variation might provide access to sources of variation that are currently neglected when short sequencing reads are aligned to a linear reference sequence [63–65].

## Conclusions

We constructed the first variation-aware reference graph for *Bos taurus* that improves read mapping accuracy over the linear reference sequence. The use of this novel reference structure facilitates accurate and unbiased sequence variant genotyping. Our results indicate that the construction of a widely-applicable bovine pan-genome graph is possible that enables accurate genome analyses for many diverged breeds.

## Methods

### Whole-genome sequencing data

We used short paired-end sequencing reads of 288 cattle from dairy (n=82 Brown Swiss – BSW, n=49 Holstein – HOL) and dual-purpose (n=49 Fleckvieh – FV, n=108 Original Braunvieh – OBV) breeds to detect variants that segregate in these populations. The average sequencing depth of the 288 cattle was 12.71-fold and it ranged from 3.49 to 70.04. Most of the sequencing data were generated previously [7, 12, 13, 66, 67]. Accession numbers for all animals are available in Additional file 3: Table S4.

We trimmed adapter sequences from the raw data and discarded reads for which the phred-scaled quality was below 15 for more than 15% of the bases using *fastp* (Chen at al., 2018). Subsequently, the sequencing reads were aligned to the linear reference assembly of the bovine genome (ARS-UCD1.2, GCF_002263795.1) using *BWA mem* [34]. Duplicates were marked and the aligned reads were coordinate sorted using the *Picard tools* software suite (http://broadinstitute.github.io/picard) and *Sambamba* [68], respectively. We discovered and genotyped polymorphic sites from the linear read alignments using the Best Practices Workflow descriptions for multi-sample variant calling with *GATK* (version 4.1.0) [1]. Because a truth set of variants required for variant quality score recalibration (VQSR) is not available for *Bos taurus*, we followed the recommendations for sequence variant discovery and filtration when applying VQSR is not possible. Genotypes of the hard-filtered variants were subsequently refined and sporadically missing genotypes were imputed with *BEAGLE* v4 [69] using the genotype likelihoods from the *GATK HaplotypeCaller* model as input values. Additional information on the sequence variant genotyping workflow and the expected genotyping accuracy can be found in Crysnanto *et al.* [12]. Nucleotide diversity was calculated in non-overlapping 10 kb windows separately for each breed using the π (nucleotide diversity) module implemented in the *vcftools* software [70].

We discovered and genotyped large structural variants (> 50 bp) including insertions, deletions, inversions, duplications and translocations in 82 sequenced BSW animals using *Delly* v0.7.8 [71] with the default settings. We retained only insertion and deletion variants that had been refined using split-reads (PRECISE-flag in the vcf file).

The principal components of a genomic relationship matrix constructed from whole-genome sequence variant genotypes were calculated using *PLINK* v1.9 [72]. The top principal components separated the animals by breeds, corroborating that the four breeds are genetically distinct (Figure 2a). To take haplotype diversity and different linkage disequilibrium phases across breeds into account, the sequence variant genotypes were phased for each breed separately using *BEAGLE* v5 [73].

Unless stated otherwise, our analyses included 541,876 biallelic SNPs and Indels that were detected on bovine chromosome 25. The *vg toolkit* version 1.17.0 “Candida” [22] was used for all graph-based analyses.

### Haplotype-aware simulation of short sequencing reads

We simulated 10 million reads (150 bp) from reference haplotypes of one animal per breed that had sequencing coverage greater than 20-fold (see Additional file 3 Table S4). Therefore, we added the phased sequence variants of each of the four animals to the linear reference to construct individualized reference graphs using *vg construct*. The haplotype-aware indexes of the resulting graphs were built using *vg index xg* and *gbwt*. *vg paths* and *vg mod* were used to extract the haplotype paths from the individualized reference graphs. Subsequently, we simulated 2.5 million paired-end reads (2×150 nt) from each haplotype using *vg sim*, yielding 10 million 150 bp reads per breed corresponding to approximately 35-fold sequencing coverage of bovine chromosome 25. The simulation parameter setting for read and fragment length was 150 and 500 (± 50), respectively. The substitution and indel error rate was 0.01 and 0.002, respectively, according to the settings used in Garrison *et al.* [22].

### Read mapping to graphs augmented with variants filtered for allele frequency

The alternate allele frequency of 541,876 variants of bovine chromosome 25 was calculated separately for the BSW, FV, HOL and OBV breeds using sequence variant genotypes of 82, 49, 49 and 108 sequenced cattle, respectively. We added to each breed-specific genome graph 20 sets of variants that were filtered for alternate allele frequency using thresholds between 0 and 1 with increments of 0.01 and 0.1 for frequency below and above 0.1, respectively. For instance, at an alternate allele frequency threshold of 0.05, the graph was constructed with variants that had alternate allele frequency greater than 5%. Alleles that were only detected in the four animals used to simulate reads (see above) were not added to the breed-specific augmented genome graphs.

The four breed-specific augmented genome graphs contained the same number of variants at a given allele frequency threshold to ensure that their density of information was similar. The number of variants added to the graphs was determined according to the breed in which the fewest variants were detected at a given allele frequency threshold. For the other three breeds, we sampled randomly from all variants that were detected at the respective alternate allele frequency threshold. We indexed the breed-specific augmented graphs using *vg index* to obtain the topological (*xg*), query (*gcsa*), and haplotype (*gbwt*) index. Eventually, the simulated reads were aligned to the breed-specific augmented reference graphs using *vg map* with default mapping parameter settings considering both graph (*xg*, *gcsa*) and haplotype (*gbwt*) indexes.

To compare the accuracy of read mapping between variation-aware and linear reference structures, the simulated reads were also aligned to the linear reference sequence of bovine chromosome 25 using either *BWA mem* with default parameter settings or *vg map*. To enable linear mapping with *vg map*, we constructed an empty graph (without adding any sequence variants) from the linear reference sequence.

### Read mapping to human population-specific augmented genome graphs

We downloaded phased whole-genome variants of 2,504 individuals from phase 3 of the 1000 Genomes Project [28] as well as the corresponding reference sequence (g1k_v37; https://www.internationalgenome.org/category/reference/). We selected four populations which we considered to be genetically distinct based on the results of a principal components analysis and for which the number of individuals was similar to the number of individuals for the four cattle breeds, i.e., GBR (British in England and Scotland, European), YRI (Yoruba in Ibadan Nigeria, African), JPT (Japanese in Tokyo, East Asia), and STU (Sri Lankan Tamil, South Asia). The principal components analysis of a genomic relationship matrix constructed using 81.27 million autosomal variants using the PLINK (v1.9) software [72]. Alternate allele frequency was calculated separately for the four populations for all variants of human chromosome 19. Nucleotide diversity was calculated with the *vcftools* software as detailed above. In order to construct population-specific augmented genome graphs, we used the reference sequence (g1k_v37) of human chromosome 19 as a backbone and added variants filtered for alternate allele frequency in the four populations (following the approach explained above). For each population, we constructed 20 graphs that contained between 3153 and 290,593 variants. We simulated 10 million paired-end reads for each population from reference haplotypes (as detailed above) of four selected samples (GBR: HG00096, YRI: NA18486, JPT: NA18939, STU: HG03642). The simulated reads were then mapped to the population-specific augmented genome graphs using the *vg toolkit*.

### Read mapping to bovine breed-specific augmented graphs

We simulated 10 million reads from the haplotypes of a BSW animal (SAMEA6272105) and mapped them to variation-aware reference graphs that were constructed using variants (SNPs and Indels) filtered for alternate allele frequency greater than 0.03. Alleles that were only detected in SAMEA6272105 were excluded from the graphs. All graphs contained 243,145 variants. The number of variants was determined according to the HOL cattle breed because the lowest number of variants segregated at an alternate allele frequency greater than 0.03 in that breed. To investigate the utility of *targeted* genome graphs, we mapped the simulated BSW reads to a graph that contained variants filtered for allele frequency in BSW cattle. To investigate *across-breed* mapping, we mapped the simulated BSW reads to graphs that contained variants filtered for allele frequency in either FV, HOL or OBV cattle. We also mapped the BSW reads to a bovine *pan-genome* graph that contained variants that were filtered for allele frequencies across the four cattle breeds. Additionally, we investigated the accuracy of mapping reads to a graph that was built from randomly selected variants. To construct the *random* graph, we randomly sampled from 2,294,416 variants that were detected on bovine chromosome 25 from animals of various breeds of cattle (http://www.1000bullgenomes.com/doco/ARS1.2PlusY_BQSR_v2.vcf.gz). The allele frequencies and haplotype phases of the random variants were not known. We constructed *personalized* graphs that contained only variants and haplotypes that were detected in the animals used for read simulation. The variation-aware graphs were subsequently indexed using *vg index* (see above). The simulated BSW reads were mapped to the different graphs using *vg map* (see above). The construction and indexing of graphs as well as read simulation and mapping was repeated ten times. We report in the main part of the paper the average values of ten replicates. This entire procedure was repeated with reads that were simulated from the haplotypes of FV (SAMN02671626), HOL (SAMN02671584) and OBV animals (SAMEA5059743).

### Read mapping to consensus reference sequences

We modified alleles of the ARS-UCD1.2 linear reference sequence using the *vcf2diploid* tool [52]. We created two adjusted linear reference sequences for bovine chromosome 25:

- *major-BSW*: 67,142 nucleotides of the linear reference sequence were replaced with the corresponding major alleles detected in 82 BSW cattle.
- *major-pan*: 73,011 nucleotides of the linear reference sequence were replaced with the corresponding major alleles detected in 288 cattle from four breeds.

Ten million BSW reads were simulated (see above) and mapped to the original and modified linear reference sequences, as well as the corresponding variation aware reference structures using either *BWA* mem or *vg map* (see above) with default parameter settings. Since the replacement of reference alleles with Indels causes a shift in the reference coordinate system, we converted the coordinates of simulated reads between the original and modified reference using a local instance of the *UCSC liftOver* tool [74] that was guided using a *chain file* produced by *vcf2diploid*. In order to prevent possible errors arising from coordinate shifts when reference nucleotides are either deleted or inserted at Indels, we repeated the analysis when only the alleles at SNPs were replaced.

### Assessment of the read mapping accuracy

We used *vg stats* to obtain the number of nodes and edges, biologically plausible paths and length for each variation-aware reference graph. To assess the accuracy of graph-based alignment, we converted the *Graph Alignment Map* (GAM)-files to JavaScript Object Notation (JSON)-files using *vg view*. Subsequently, we applied the command-line JSON processor *jq* (https://stedolan.github.io/jq/) to extract mapping information for each read. Mapping information from linear alignments were extracted from the *Binary Alignment Map* (BAM)-files using the Python module *pysam* (version 0.15.3) (https://github.com/pysam-developers/pysam).

Using *vg annotate*, we annotated the simulated reads with respect to the linear reference coordinates and determined if they contained non-reference alleles. Comparing the true and mapped positions of the simulated reads enabled us to differentiate between correctly and incorrectly mapped reads. Following the approach of Garrison *et al.* [22] and taking into account the possibility that aligned reads may be clipped at Indels, we considered reads as incorrectly mapped if their starting positions were more than *k=150* (k=read length) bases distant from true positions. The functional relevance genomic regions where the simulated reads originated from were determined based on the Ensembl annotation (version 99, [75]) of the bovine ARS-UCD 1.2 reference sequence. The coordinates of repetitive elements were determined based on RepeatMasker [76] annotation tables of the UCSC Genome Browser.

In order to assess mapping sensitivity and specificity, we calculated the cumulative TPR (*True positive rate*) and FPR (*False positive rate*) at different mapping quality thresholds and visualized it as pseudo-ROC (*Receiver Operating Characteristic*) curve [22] using:

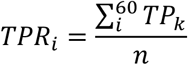

and

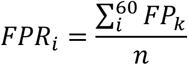

*where* TP_i_ and FP_i_ represent the number of correctly and incorrectly mapped reads, respectively, at a given phred-scaled mapping quality threshold i (60, 50, 40, 30, 20, 10, 0), and n is the total number of reads mapped.

### Read mapping and sequence variant genotyping from bovine whole-genome graph

Using 14,163,824 autosomal biallelic variants (12,765,895 SNPs and 1,397,929 Indels) that had alternate allele frequency greater than 0.03 in 82 BSW cattle, we constructed a BSW-specific augmented whole-genome graph. The Hereford-based linear reference sequence (ARS-UCD1.2) was the backbone of the graph. Specifically, we constructed graphs for each of the 29 autosomes separately using *vg construct*. Subsequently, *vg ids* was run to ensure that the node identifiers were unique in the concatenated whole-genome graph. We removed complex regions from the whole-genome graph using *vg prune* with default parameter settings and built the topological (*xg*) and query (*gcsa*) index for the full and pruned graph, respectively, using *vg index.* The haplotype paths of the 82 BSW cattle obtained using *BEAGLE* v5 (see above) were provided using a *gbwt* index.

To evaluate sequence variant genotyping from the whole-genome graph, we used between 122,753,846 and 904,047,450 million paired-end (2×150 bp) sequencing reads from 10 BSW cattle (SAMEA6163185, SAMEA6163188, SAMEA6163187, SAMEA6163177, SAMEA6163178, SAMEA6163176, SAMEA6163179, SAMEA6163183, SAMEA6163181, SAMEA6163182, Additional file 3 Table S4) that had been sequenced at between 5.74 and 39.88-fold genome coverage. These animals were not part of the 82 BSW animals that were used to detect the variants that were added to the graph. We trimmed adapter sequences and removed reads that had more than 20% bases with phred-scaled quality less than 20 using *fastp* [77]. Subsequently, we mapped the pruned reads to either the BSW-specific augmented whole-genome graph or the linear reference sequence using either *vg map* while supplying both graph (*xg*, *gcsa*) and haplotype (*gbwt*) index to produce GAM files for each sample or *BWA mem*. To make the coordinates of the graph-based alignments compatible with linear reference coordinates, we converted the GAM- to BAM-files using *vg surject*. Variants were detected and genotyped from the surjected files using the multi-sample variant calling approach of either *GATK* [39], *Graphtyper* [40], or *SAMtools* [38], as stated above and detailed in [12].

In order to assess the read mapping accuracy from real sequencing data, we calculated the proportion of reads that aligned (i) perfectly and (ii) uniquely [18, 35, 78]. A read was considered to map perfectly if the edit distance was zero along the entire read (NM:0 tag in *BWA* mem-aligned BAM files; identity 1 in *vg map*-aligned GAM-files), and without hard clipping (H tag) or soft clipping (S tag) in CIGAR string. A read was considered to map uniquely if either a single primary alignment was reported for the respective read or reads that had secondary alignments (XA tag in *BWA mem*-aligned BAM files; secondary_score > 0 in *vg map*-aligned GAM-files) had one alignment with phred-scaled mapping quality score of 60.

The sequenced BSW animals also had been genotyped at between 24,512 and 683,752 positions using Illumina SNP Bead chips. The sequence variant genotypes were compared to microarray-called genotypes at corresponding positions to calculate recall/non-reference sensitivity, genotype concordance, precision and non-reference discrepancy [1, 79]. The concordance metrics are explained in Additional file 1: Figure S17.

Snakemake workflows [80] for whole-genome graph construction, read mapping and variant discovery are available in the Github repository (https://github.com/danangcrysnanto/bovine-graphs-mapping).

### Assessment of reference allele bias

Reference allele bias was assessed at the heterozygous genotypes that had been detected in a BSW animal (SAMEA6163185) that had been sequenced at high (40-fold) coverage. Raw sequencing data were filtered as stated above and aligned to either the linear reference sequence or BSW-specific augmented genome graph using *BWA mem* and *vg map*, respectively. Sequence variant genotypes were discovered and genotyped from either surjected graph-based or linear alignments using the single sample variant calling approaches implemented in either *GATK HaplotypeCaller* or *SAMtools mpileup*. Variants were filtered using quality by depth (QD) >10, mapping quality (MQ) > 40 and minimum read depth (DP) greater than 25 to ensure confident genotype calls and sufficient support for reference and alternate alleles at heterozygous genotypes. We considered only variants that were detected from both graph-based and linear alignments. At each heterozygous genotype, we quantified the number of reads supporting alternate and reference alleles using allelic depth information from the vcf files.

## Declarations

### Ethics approval and consent to participate

Not applicable.

### Consent for publication

Not applicable.

### Availability of data and materials

The scripts and data used in this study are available via *GitHub* repository (https://github.com/danangcrysnanto/bovine-graphs-mapping) and archived in *Zenodo* (data: https://doi.org/10.5281/zenodo.3759712 [46] and scripts: https://doi.org/10.5281/zenodo.3763286 [81]). Raw sequencing read data of 298 cattle used for graph construction, evaluation of variant genotyping accuracy and assessment of reference allele bias are available at the European Nucleotide Archive (ENA) (http://www.ebi.ac.uk/ena). Accession numbers are provided in Additional file 3 Table S4.

### Author contributions

Analyzed data: DC HP; Wrote the paper: DC HP; Read and approved the final version of the paper: DC HP.

## Acknowledgements

We thank the many people who made data and software publicly available, particularly *vgteam* for providing the *vg toolkit* as an open source software. We thank Braunvieh Schweiz for providing pedigree and genotype data of Original Braunvieh and Brown Swiss cattle. Semen samples of the sequenced bulls were kindly provided by Swissgenetics.

## Additional files

### Additional file 1

File format: pdf

Description: Integrated supplementary figures: S1-S17.

### Additional file 2

File format: pdf

Description: Integrated supplementary notes: S1-S3.

### Additional file 3

File format: xlsx

Description: Integrated supplementary tables: S1-S4.

