## Additional file 1 for "Bovine breed-specific augmented reference graphs facilitate accurate sequence read mapping and unbiased variant discovery"

Danang Crysnanto and Hubert Pausch
Animal Genomics, ETH Zürich, Zürich, Switzerland

### Integrated supplementary figures

**Figure S1: Number of 256 bp haplotype paths in the graphs with an increasing number of variants added to the graphs.**
The line plot is fitted using *loess* function in R.

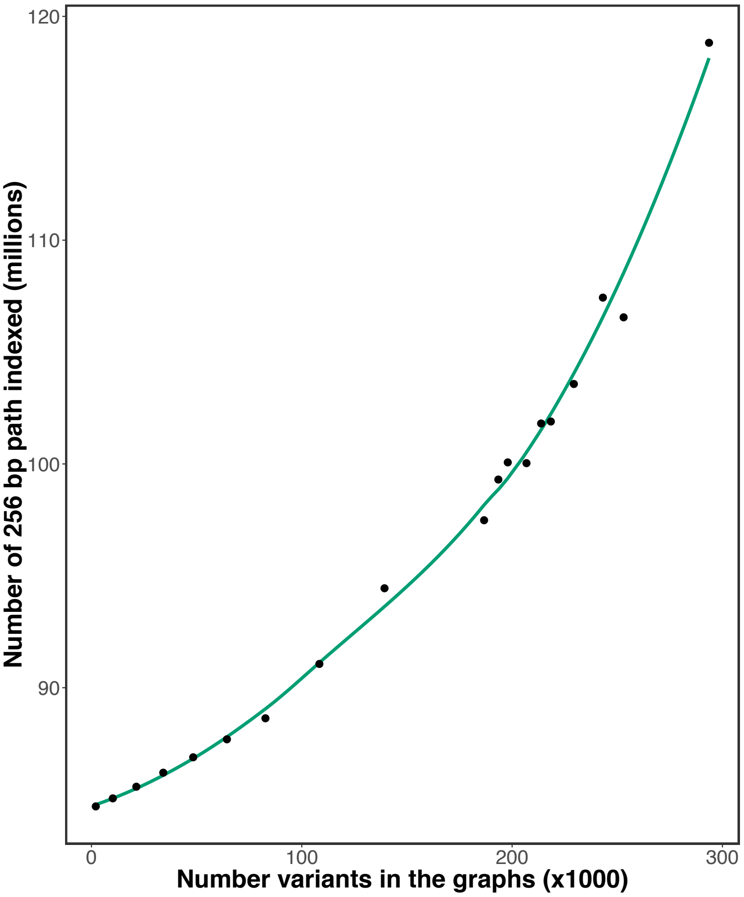

**Figure S2: Single-end mapping accuracy using genome graphs that contained variants filtered for allele frequency.**
(a) Proportion of incorrectly mapped reads for four breed-specific augmented genome graphs. Diamonds and large dots represent results from linear mapping using *BWA mem* and *vg*, respectively. The inset is a larger representation of the mapping accuracy for alternate allele frequency thresholds less than 0.1. (b) Read mapping accuracy for breed-specific augmented graphs that contained variants that were either filtered for alternate allele frequency (triangles) or sampled randomly (circles) from all variants detected within a breed. The dashed and solid line represents the average proportion of mapping errors across four breeds using variant prioritization and random sampling.

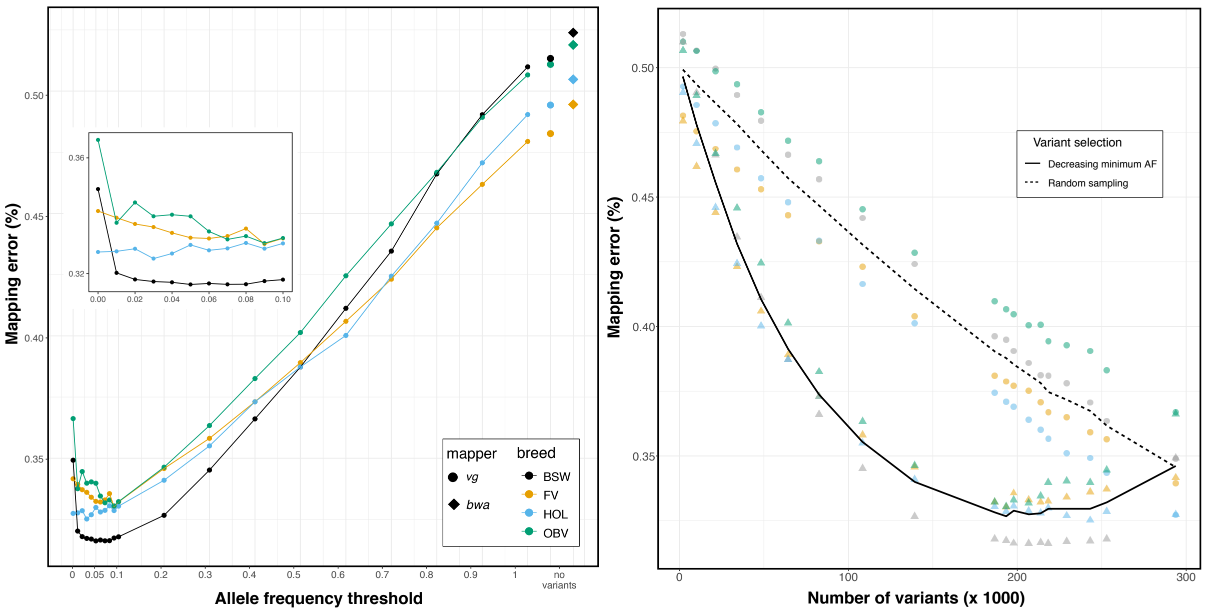

**Figure S3: Number of variants detected on chromosome 25 in 82 BSW, 49 FV, 49 HOL and 108 OBV cattle.**
Variants are binned according to allele frequency.

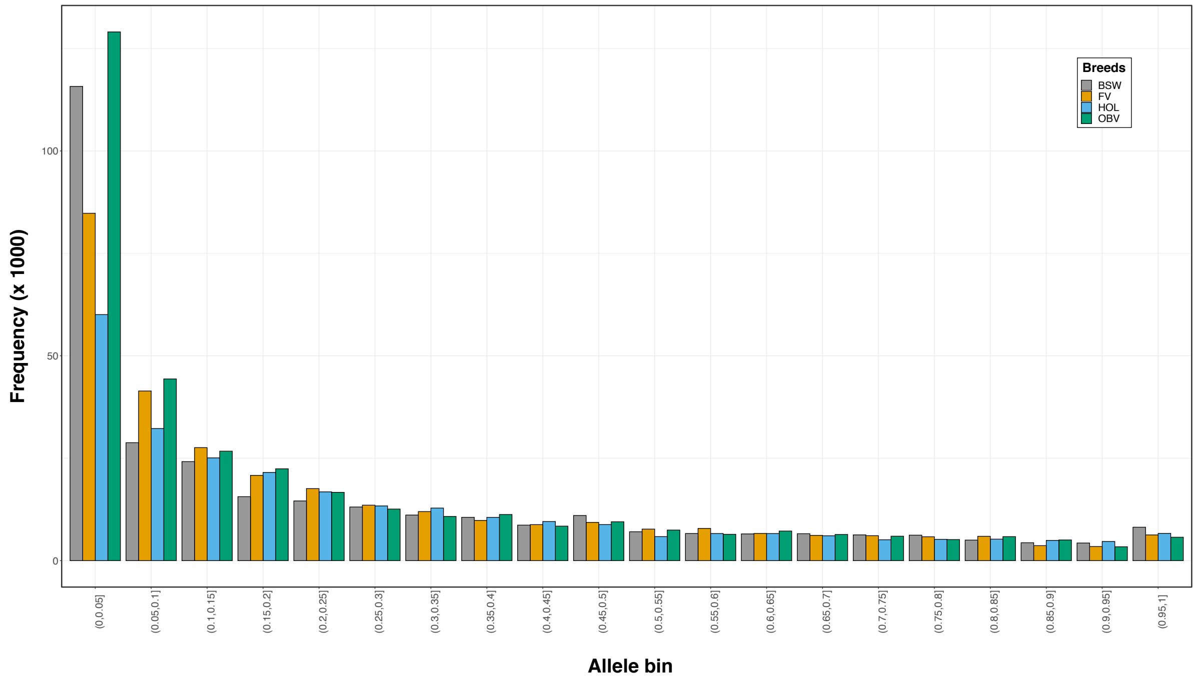

**Figure S4: Distribution of alternate allele frequencies in four cattle breeds and four human populations based on (a) bta25 and human chromosome 19 used for graph construction, and (b) whole genome variants.**
The bars indicate the proportion of sequence variants for 20 allele frequency classes. Different colour indicates cattle breeds (HOL, FV, BSW, OBV) and human populations (JPT, GBR, STU, YRI).

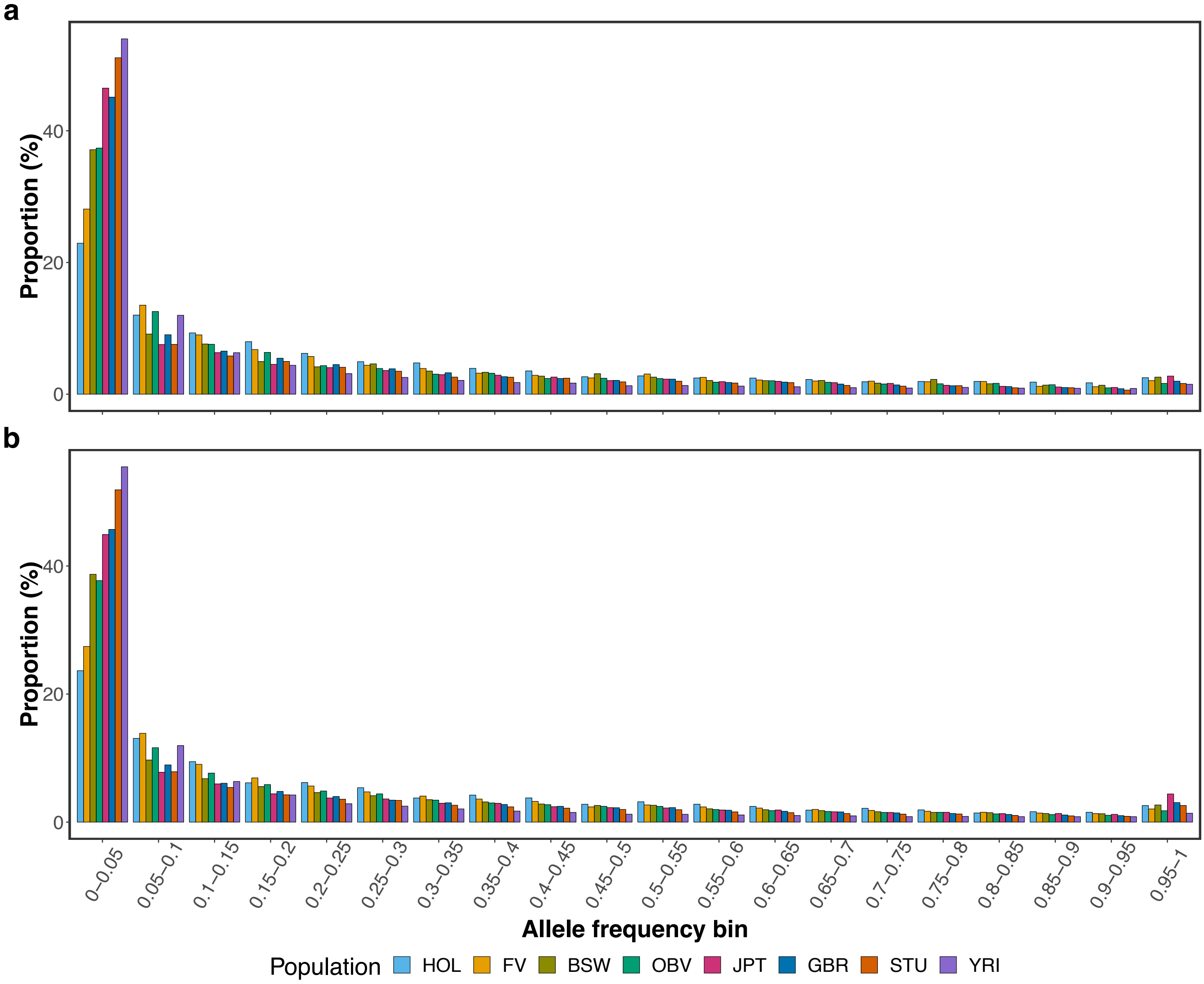

**Figure S5: Nucleotide diversity (π) based on whole genome autosomal variants in cattle and human.**

Nucleotide diversity (π) from each population calculated using vcftools with 10 kb non-overlapped windows based on whole genome autosomal variants. Number under the boxplot indicates average across windows.

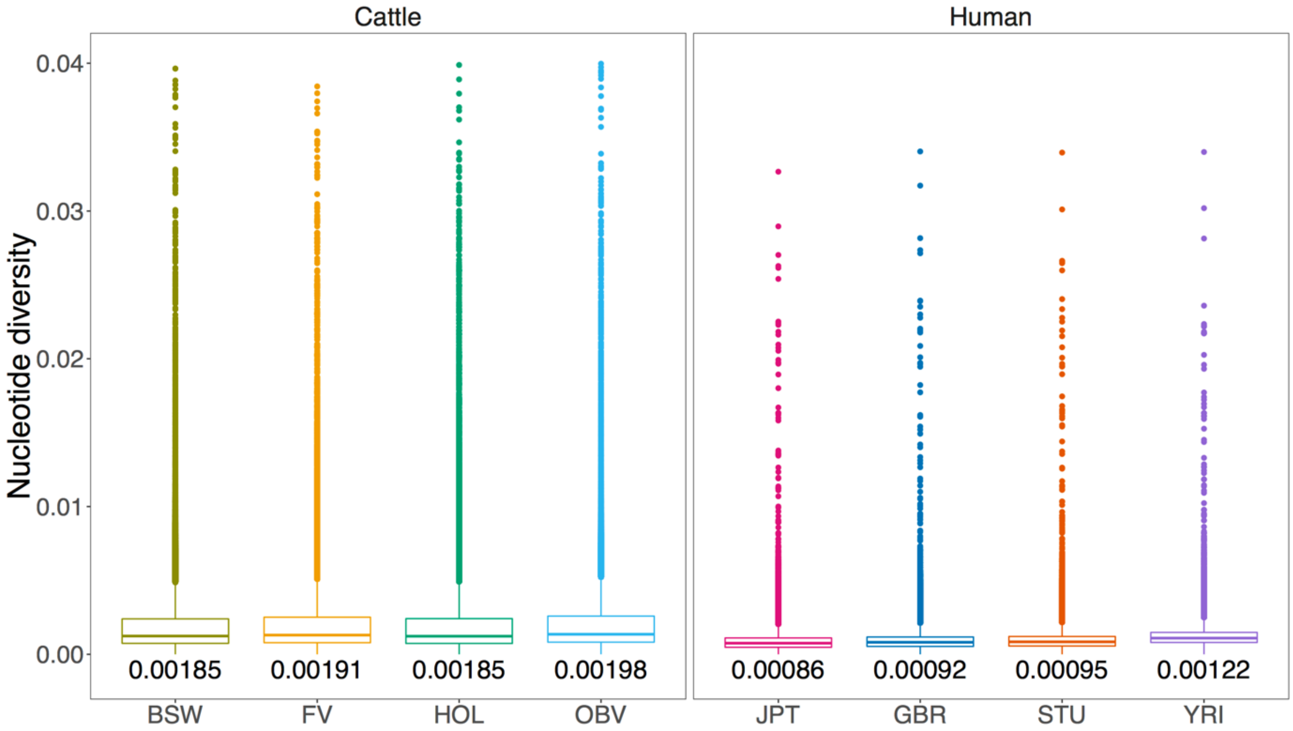

**Figure S6: Single-end mapping accuracy using four human population-specific augmented graphs.** (a) Proportion of incorrectly mapped reads for four population-specific augmented genome graphs (b) True positive (sensitivity) and false positive mapping rate (specificity) parameterized based on the mapping quality for the best performing graph from each population. (c) Read mapping accuracy for population-specific augmented graphs that contained variants that were either filtered for alternate allele frequency (triangles) or sampled randomly (circles) from all variants detected within a population. The dashed and solid line represents the average proportion of mapping errors across four populations using variant prioritization and random sampling.

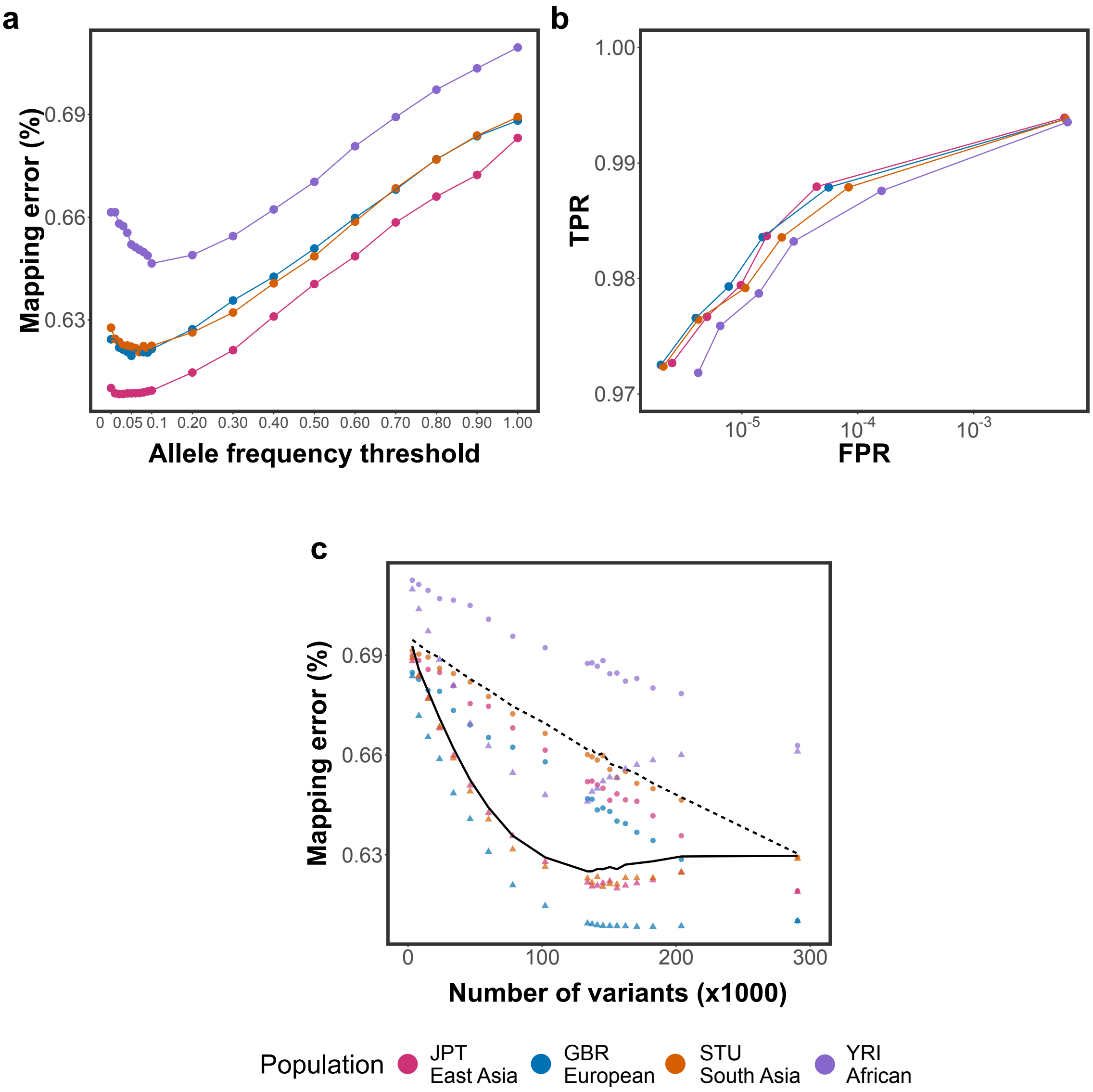

#### Figure S7: The accuracy of mapping simulated BSW single-end reads to variation-aware and linear reference structures.

(a) Proportion of BSW single-end reads that mapped erroneously against breed-specific augmented graphs, random graphs or linear reference sequences. Dark and light blue colours represent the proportion of incorrectly mapped reads with mapping quality (MQ)<10 and MQ>10, respectively. (b) True positive (sensitivity) and false positive mapping rate (specificity) parameterized based on the mapping quality. (c) Dark and light green colours represent the proportion of incorrectly mapped reads that matched corresponding reference nucleotides and contained non-reference alleles, respectively.

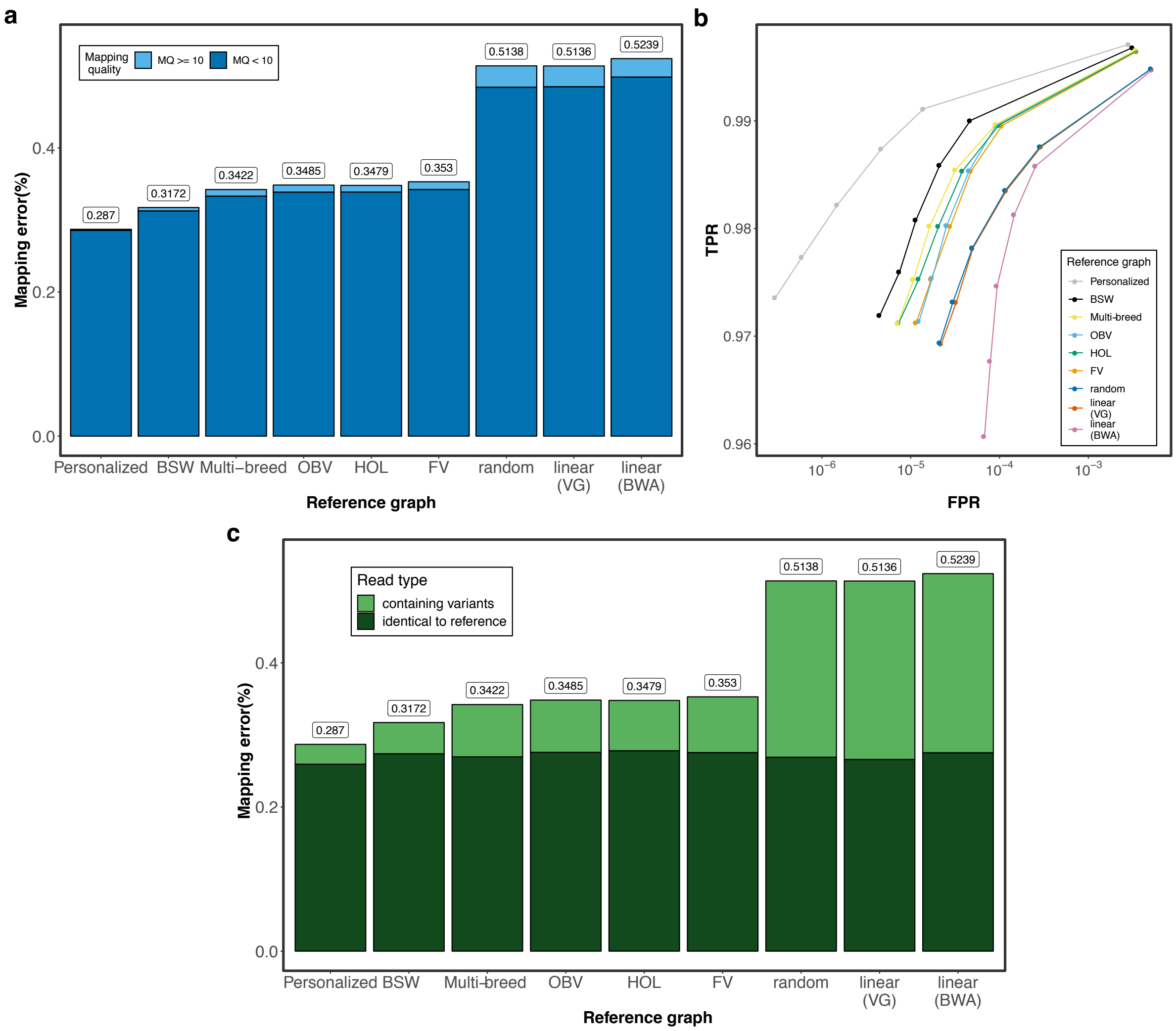

**Figure S8: Overlap of the variants** (N=243,145) between the BSW-and all other variation-aware reference graphs. The values are averaged across 10 replicates.

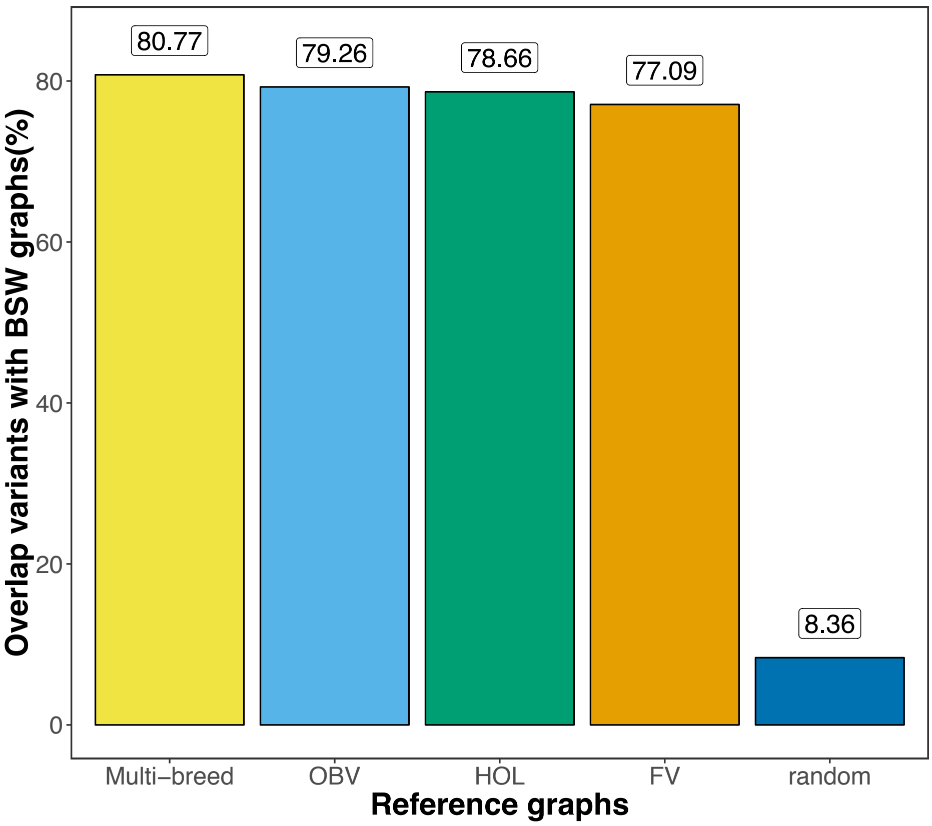

**Figure S9: Pairwise heatmap of P values from t tests** comparing 8 graph-based mapping scenarios for **(a)** paired- and **(b)** single-end reads. The P-values are adjusted for multiple testing using Bonferroni-correction.

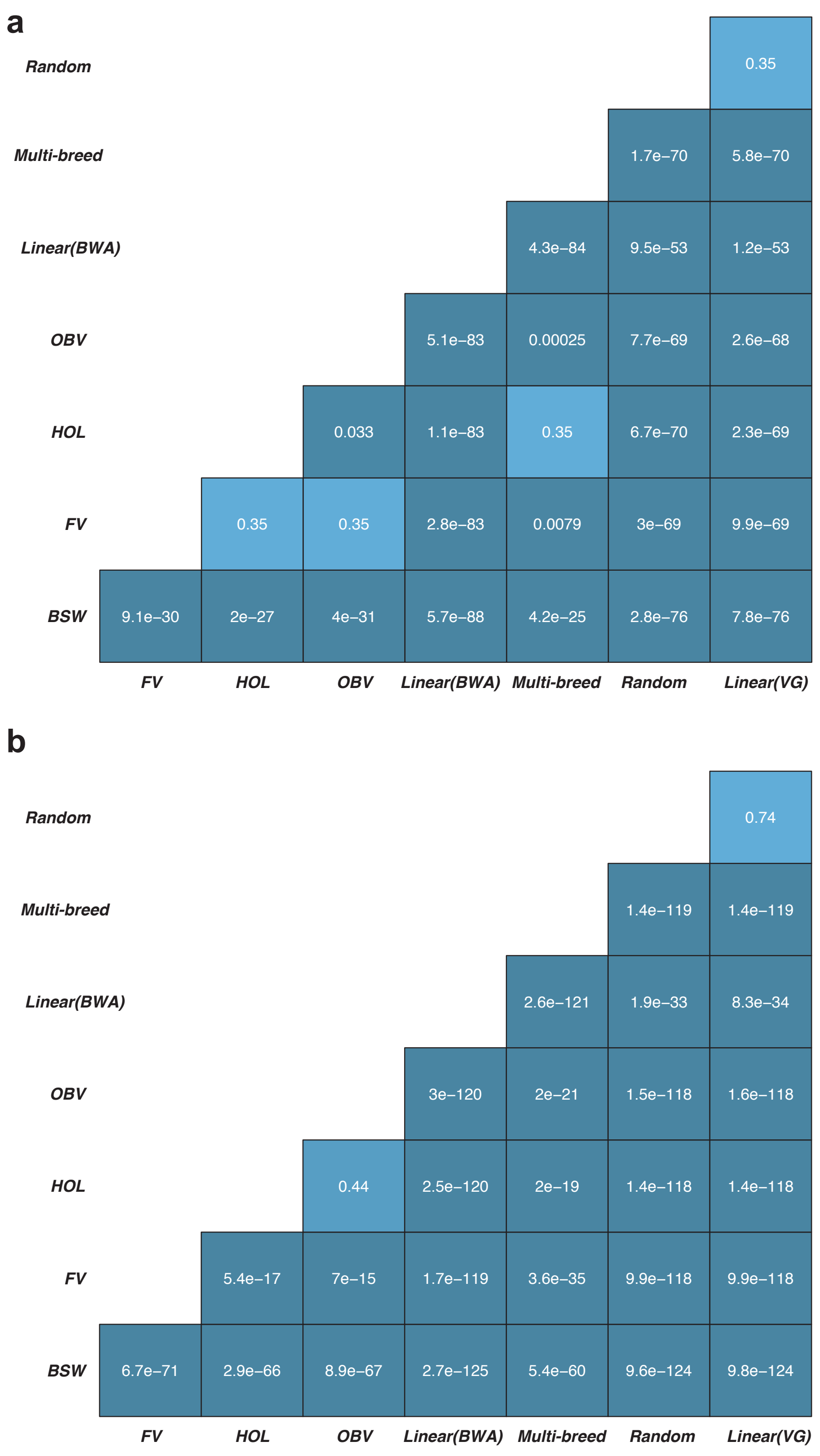

**Figure S10: The accuracy of mapping simulated FV, HOL and OBV reads to variation-aware and linear reference structures.** (a) Proportion of reads that mapped erroneously against personalized graphs, breed-specific augmented graphs, random graphs or linear reference sequences. Dark and light blue colours represent the proportion of incorrectly mapped reads with mapping quality (MQ)<10 and MQ>10, respectively. The upper and lower panels reflect paired-end and single-end reads, respectively. (b) Dark and light green colours represent the proportion of incorrectly mapped reads that matched corresponding reference nucleotides and contained non-reference alleles, respectively. The upper and lower panels reflect paired-end and single-end reads, respectively

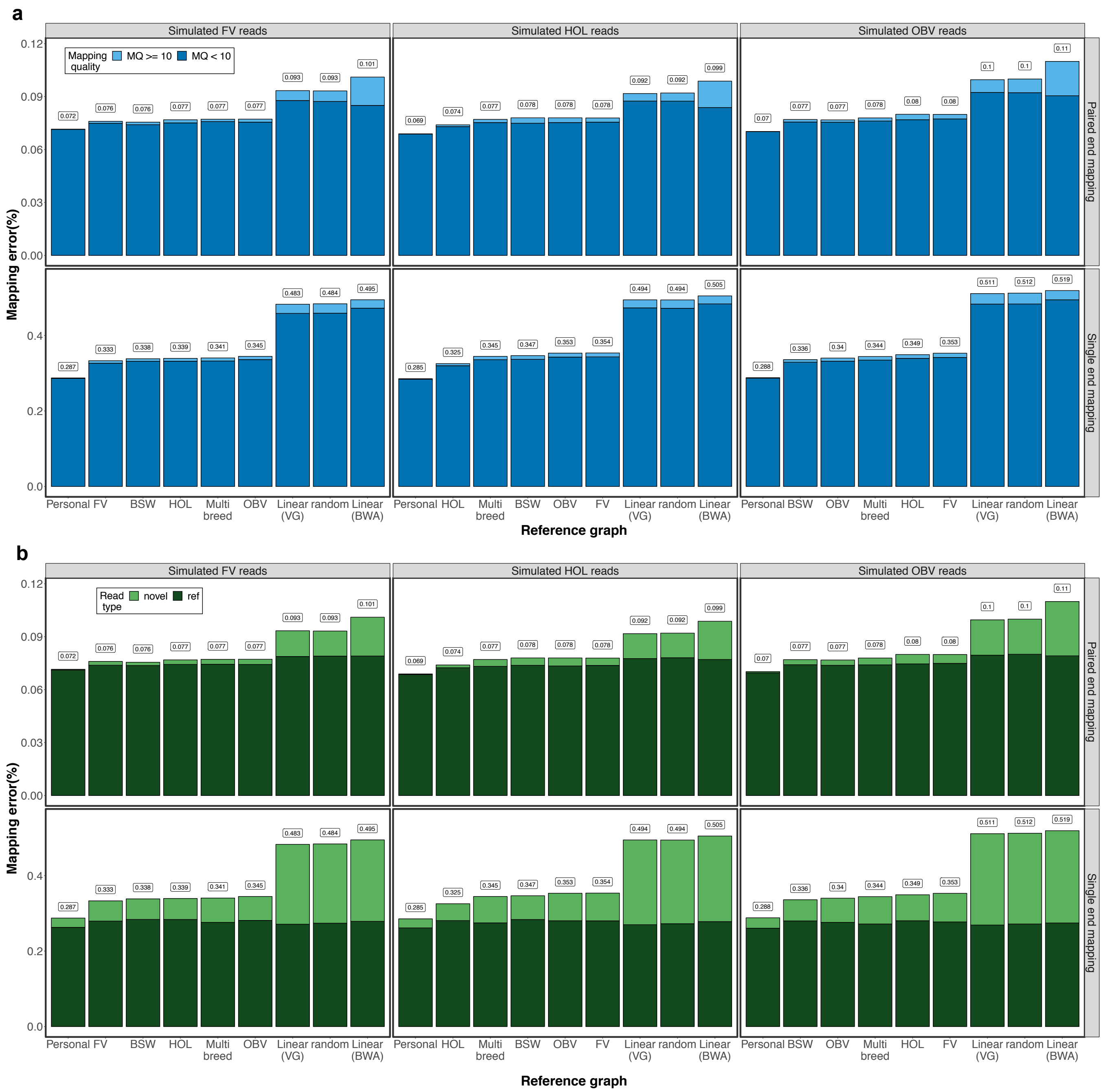

**Figure S11: ROC curves split by read's novelty**

Cumulative *True positive* and *False positive rate* at different mapping quality thresholds visualized as Receiver Operating Characteristic (ROC) curves for reads than contain variants and match corresponding reference alleles. The upper and lower panels represent results from paired- and single-end reads.

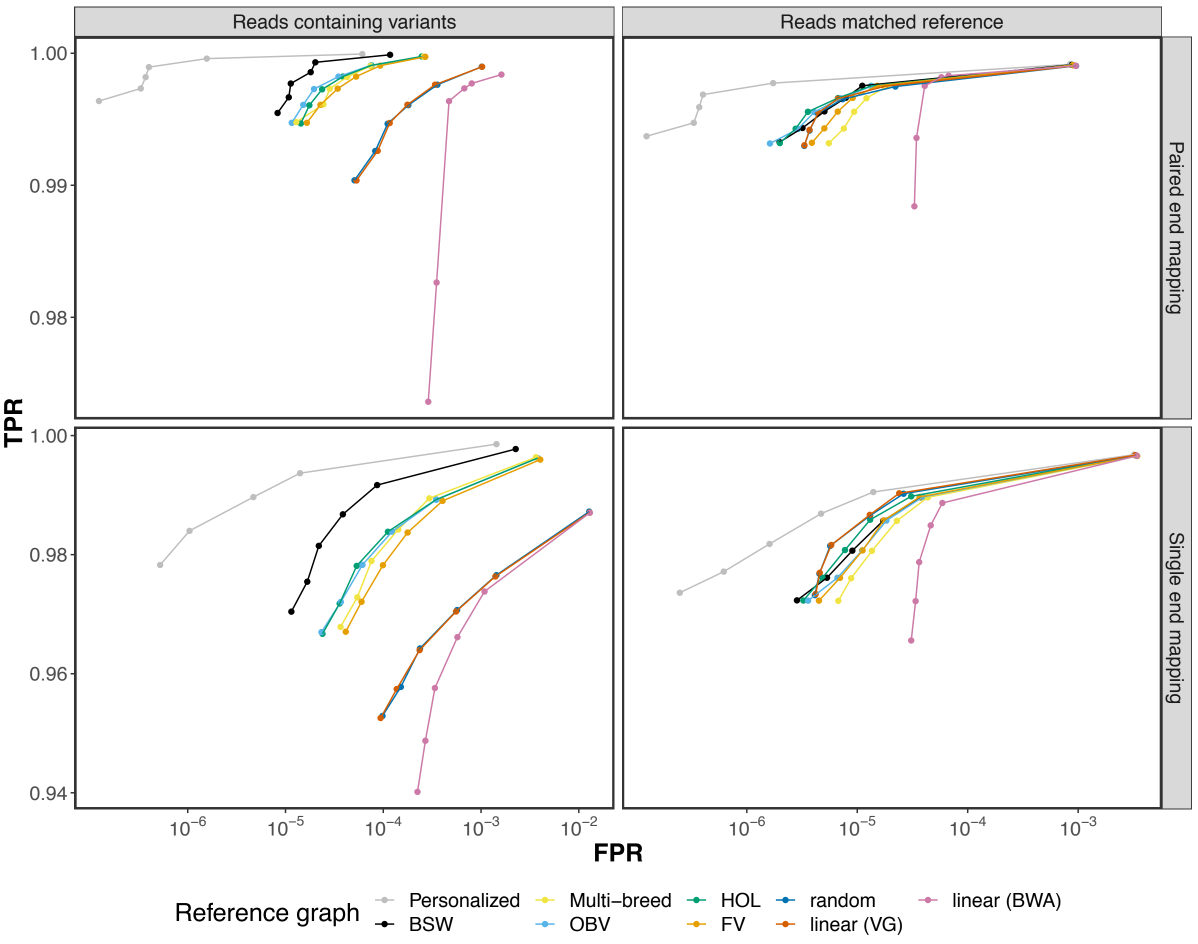

**Figure S12: Mapping accuracy for reads originating from different genomic features.**
The origin of 10 million simulated reads was determined based on the *Bos taurus* ARS-UCD1.2 ensembl 99 annotations (exonic and genic) and the ARS-UCD1.2 repeat regions labelled by Repeat Masker (Interspersed duplications including SINEs, LINEs, LTR, and DNA transposable elements, and simple repeats which contain low-complexity and simple repetitive regions). Different colour indicates the proportion of erroneously mapped reads for each annotation category. The orange bars represent the average proportion of mis-mapped reads for six graph-based (BSW, Multi-breed, OBV, HOL, FV, random) and two linear (VG, BWA) reference structures. Reads were simulated from haplotypes of a BSW individual.

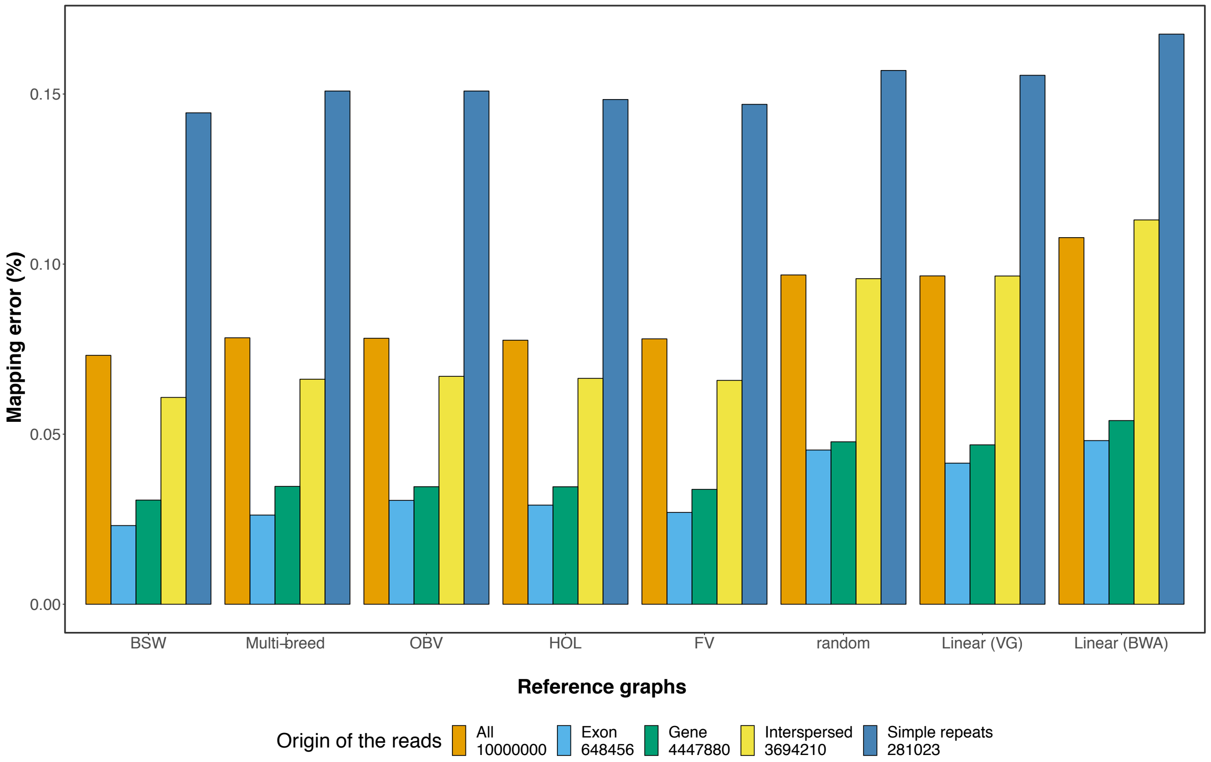

**Figure S13: Single-end read mapping accuracy using breed-specific augmented genome graphs and consensus linear reference sequences.**
(a) Dark and light blue represent the proportion of reads that mapped incorrectly using *BWA mem* and *vg*, respectively, to the BSW-specific augmented reference graph (BSW-graph), the BSW-specific (major-BSW) and multi-breed linear consensus sequence (major-pan) and the bovine linear reference sequence (unmodified). (b) True positive (sensitivity) and false positive mapping rate (specificity) parameterized based on the mapping quality. (c) Paired- and single-end read mapping accuracy using breed-specific augmented genome graphs and consensus linear reference sequences that were only adjusted at SNPs.

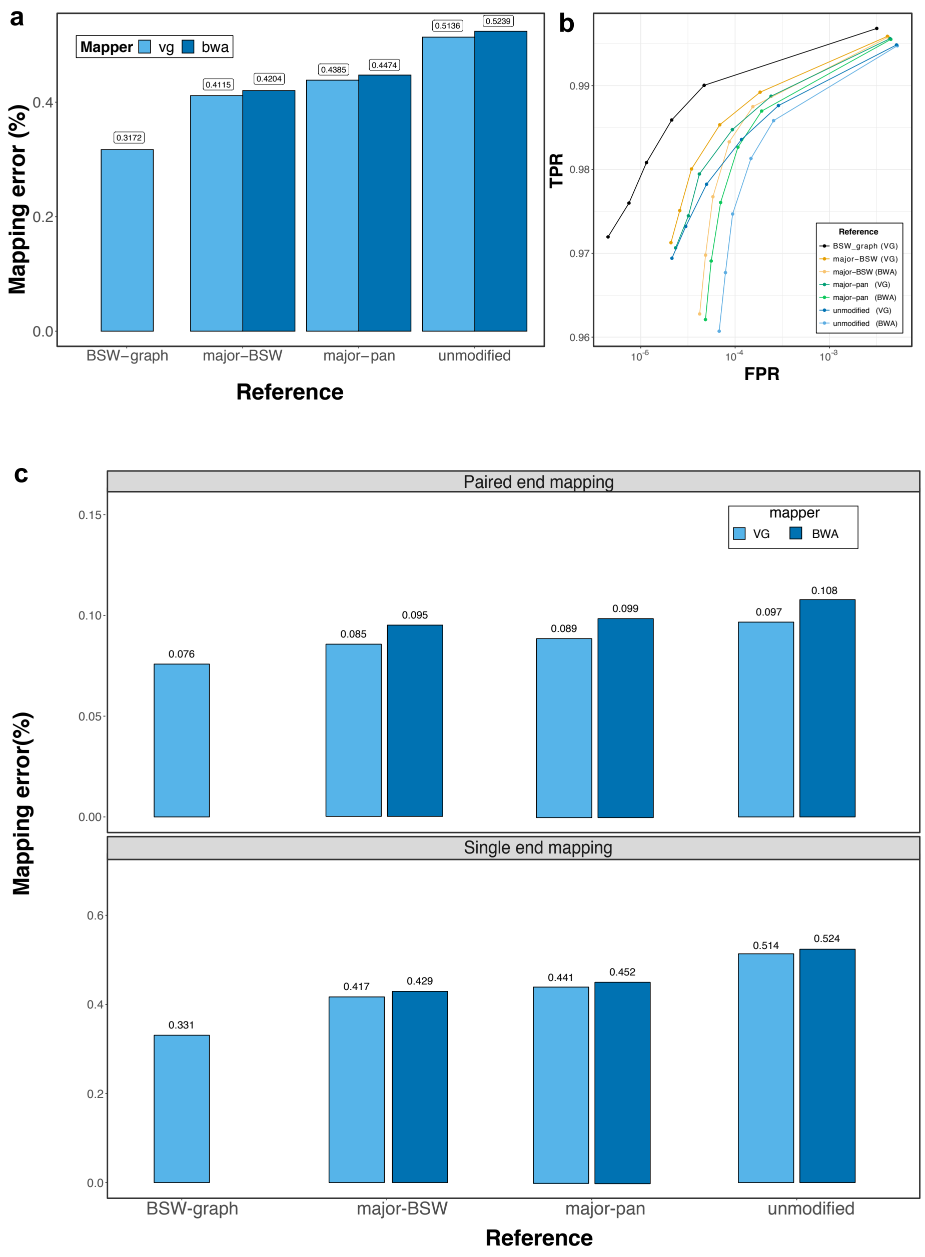

**Figure S14: Graph alignment visualization.** Visualization of a 23-bp insertion at Chr10: 5,941,270 in graph and linear alignments using the *sequence tube map* tool (Beyer et al., 2019). The variant was called heterozygous from the linear alignment, but the allelic ratio was highly biased towards the reference allele. Visual inspection suggests that more reads supporting the alternate allele are present in the graph alignments. Red and blue colour indicates forward and reverse reads, respectively. The reads from the linear alignment were realigned to the variation-aware graph for the purpose of the visualisation.

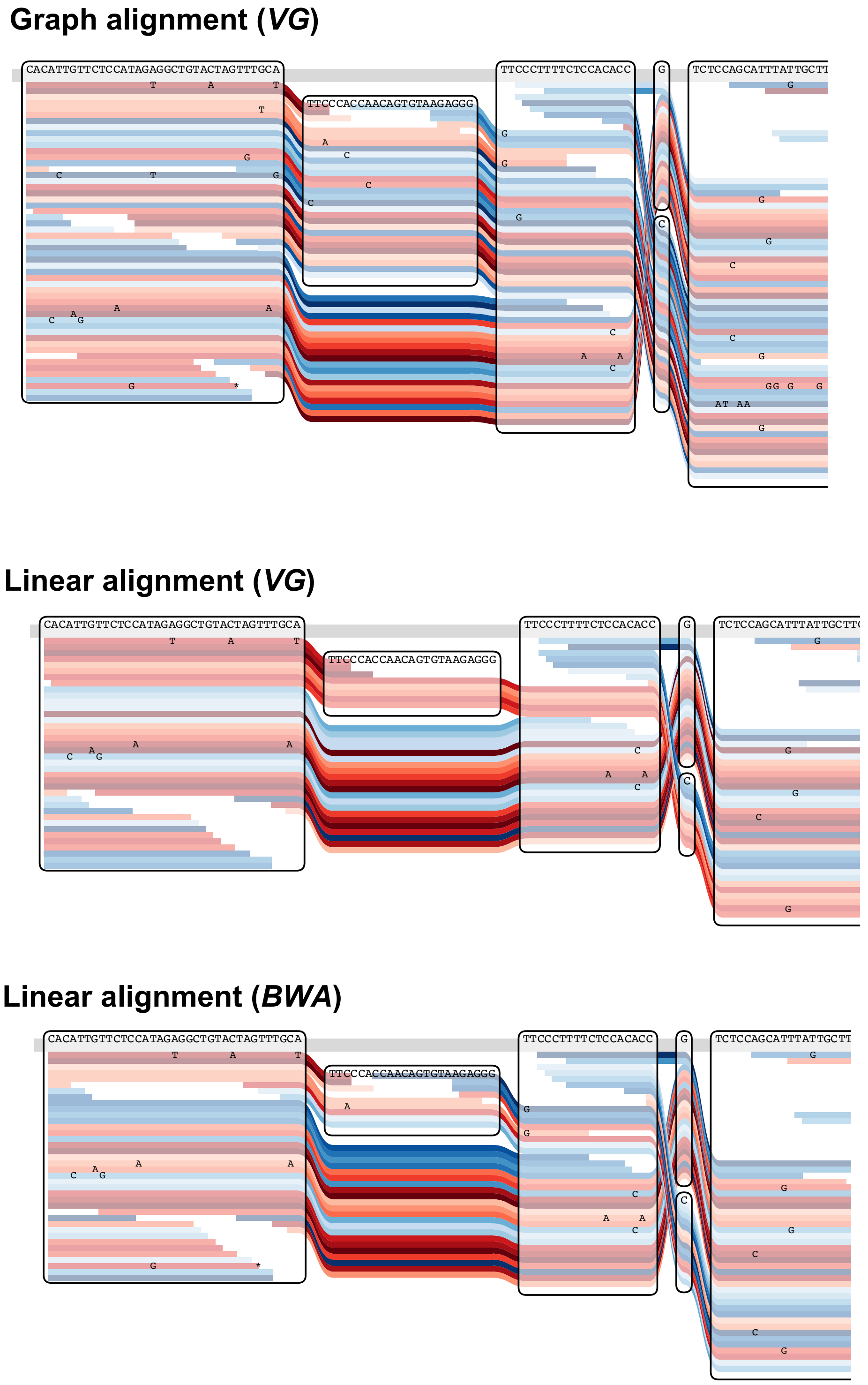

**Figure S15: Difference in the total of mapped reads, and reads support for reference and alternate alleles** between the graph-based and *BWA* alignments for deletions, SNPs and insertions. Positive values indicate a larger number of reads for graph-based alignments. The dashed grey line indicates equal support for graph-based and linear alignments. The circles represent the mean (± standard error of mean) values at a given variant length. Red and green colour indicates that the alternate allele is included and not included in the graph, respectively.

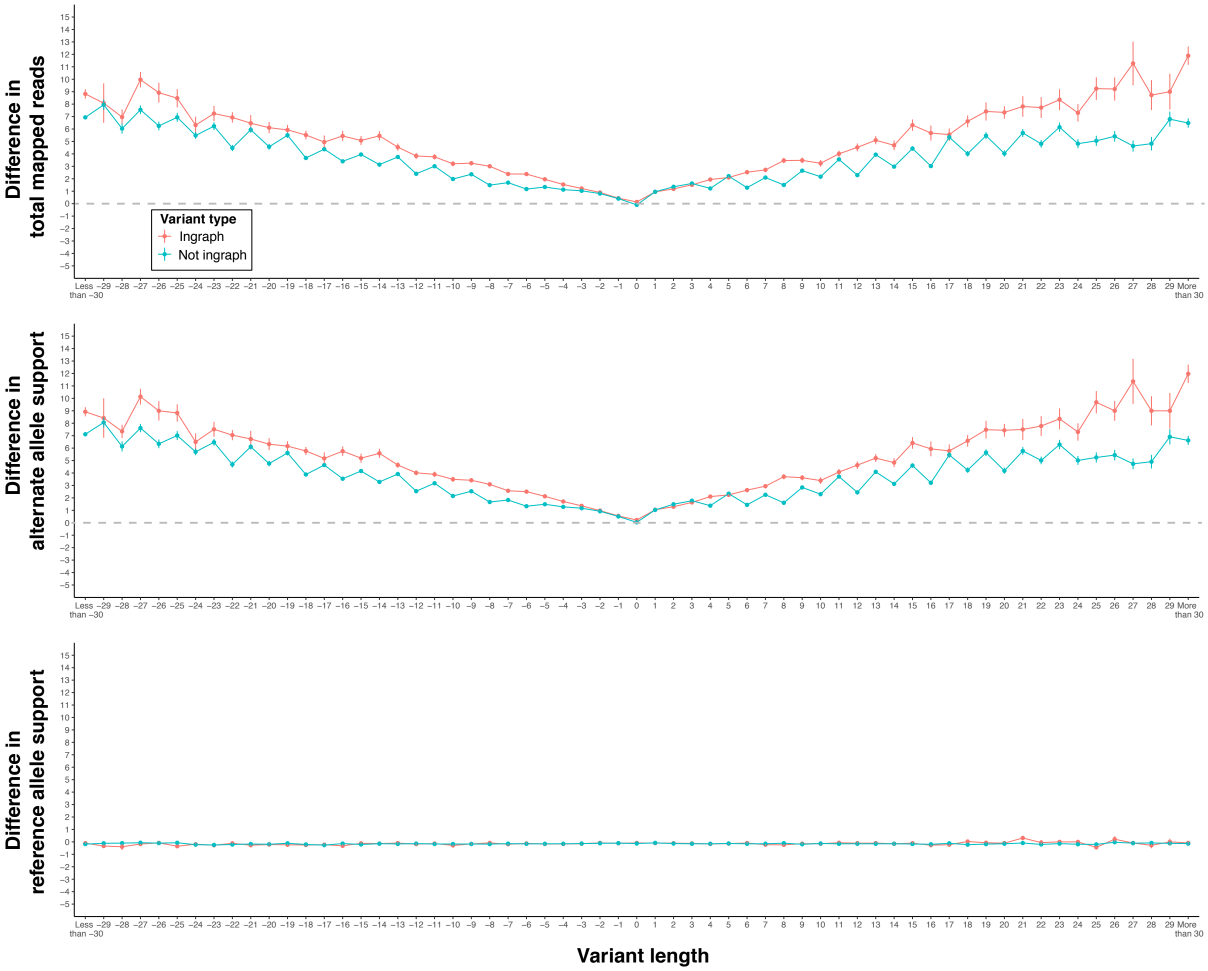

**Figure S16: Proportion of soft-clipped reads at heterozygous sites in graph (*vg*) and linear (*vg* and *BWA*) alignments.** We considered only variants for which the alternate allele was already included in the graph. The circles represent the mean (± standard error of mean) values at a given variant length.

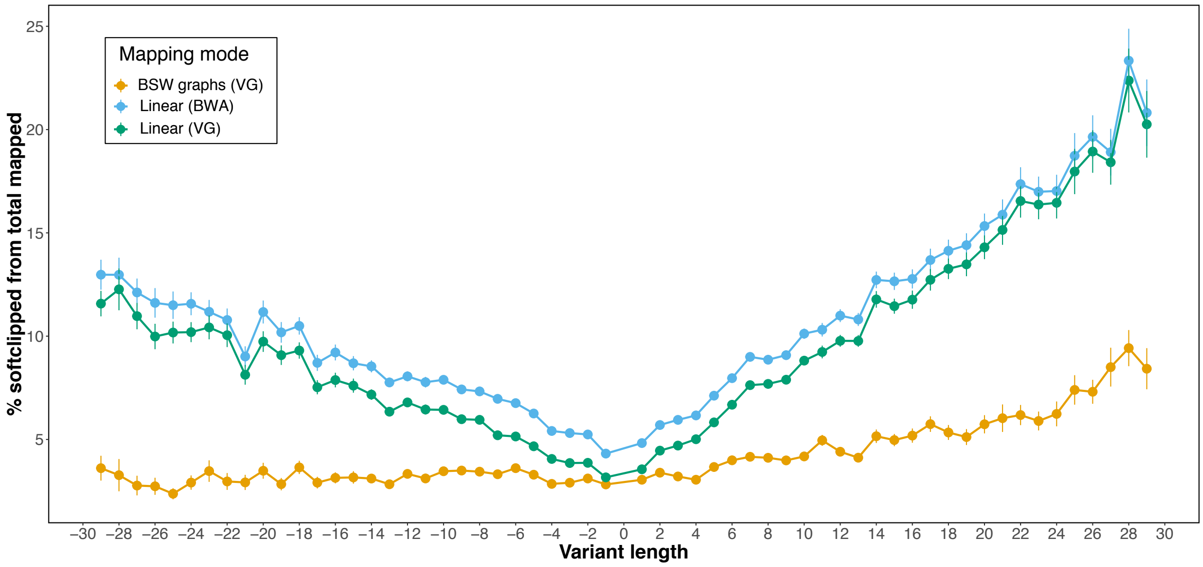

**Figure S17: Genotype concordance matrices for four quality parameters**. For each metric, we divided the sum of the red cells by the sum of the cells within the green frame.

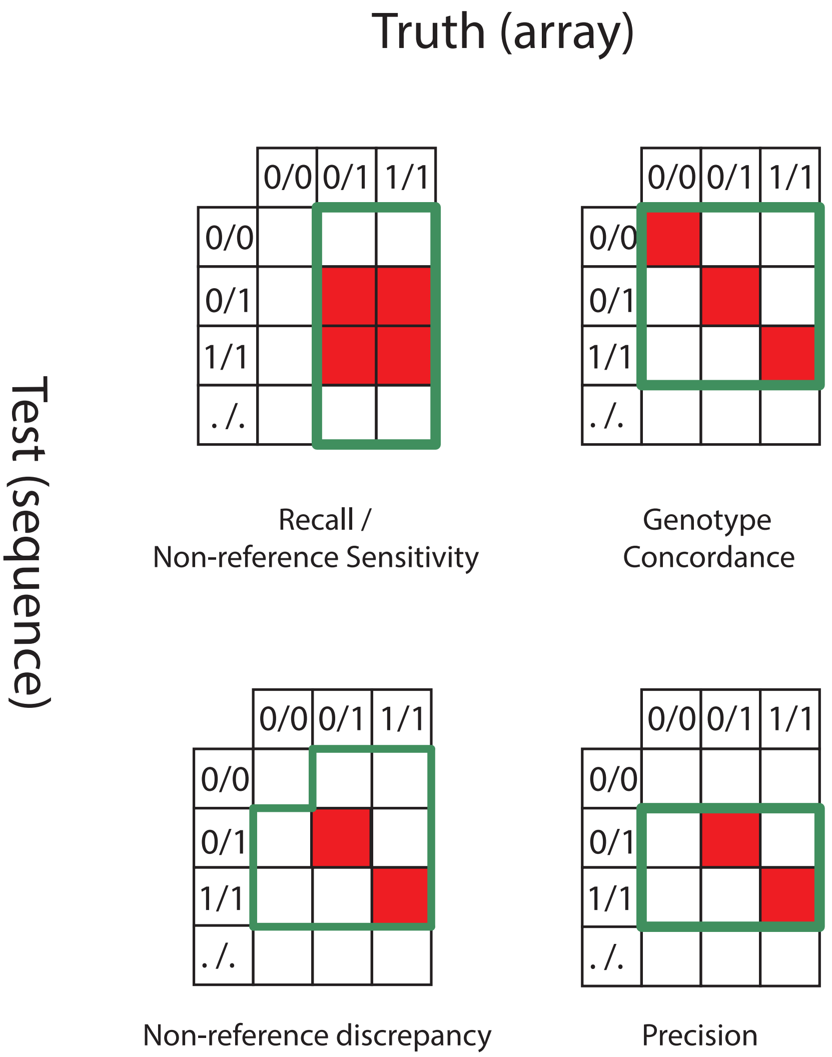
