## Additional file 2 for "Bovine breed-specific augmented reference graphs facilitate accurate sequence read mapping and unbiased variant discovery"

### Supplementary Note 1

### Comparison of variant prioritization approaches

We applied FORGe (Pritt *et al.* 2018) to prioritize variants to be added to the Brown Swiss reference graph for chromosome 25. Specifically, we considered the four variant ranking approaches implemented in FORGe and compared the mapping accuracy from the resulting graphs with a graph that was constructed with variants selected based on an allele frequency threshold.
The following prioritization approaches were investigated:

1. Pop Cov: variants ranked based on allele frequency
2. Pop Cov + blowup: variants ranked based on allele frequency and proximity (variants that are nearby receive lower scores)
3. Hybrid: variants ranked based on allele frequency and how the variants affect the resulting k-mer profile of the genome graph (variants that would increase the repetiveness of the resulting graph receive lower scores)
4. Hybrid + blowup: hybrid methods + considering variant proximity
5. AF threshold: variants ranked based on allele frequency (AF, as applied in our paper).

We refer to the FORGe paper (Pritt *et al.* 2018) for a detailed description on the implementation of the variant prioritization methods 1-4.

For each prioritization approach, we constructed a number of graphs that included the top x% of the ranked variants, where x ranged from 1 to 100 with steps of 10 (e.g., a graph constructed with x=10 included 34,715 out of 347,147 bta25 Brown Swiss variants). We then mapped paired-end reads simulated form a Brown Swiss animal (as detailed in the Material and Methods part of the main manuscript) to the graphs in order to calculate mapping accuracy.

Graphs constructed with variants that were prioritized solely using allele frequency (as applied in our current paper and the *Pop Cov* method of FORGe) enable the most accurate mapping of reads (Table SN1 & Figure SN1). Considering additional factors other than allele frequency did not lead to further accuracy improvements. The mapping accuracy of the Pop Cov and AF threshold strategies was virtually identical when the same number of variants was used. The most accurate Pop Cov approach corresponds to an alternate allele frequency threshold of 0.06.

#### Table SN1: Comparison of the most accurate graph from each ranking method

| Ranking method | Mapping error (%) | Number of variants added to the graph with maximum accuracy |
|---|---|---|
| **Pop Cov** | 0.0722 | 208288 |
| **Pop Cov + blowup** | 0.0730 | 208288 |
| **AF threshold** | 0.0723 | 208288 |
| **Hybrid** | 0.0749 | 347147 |
| **Hybrid + blowup** | 0.0749 | 347147 |

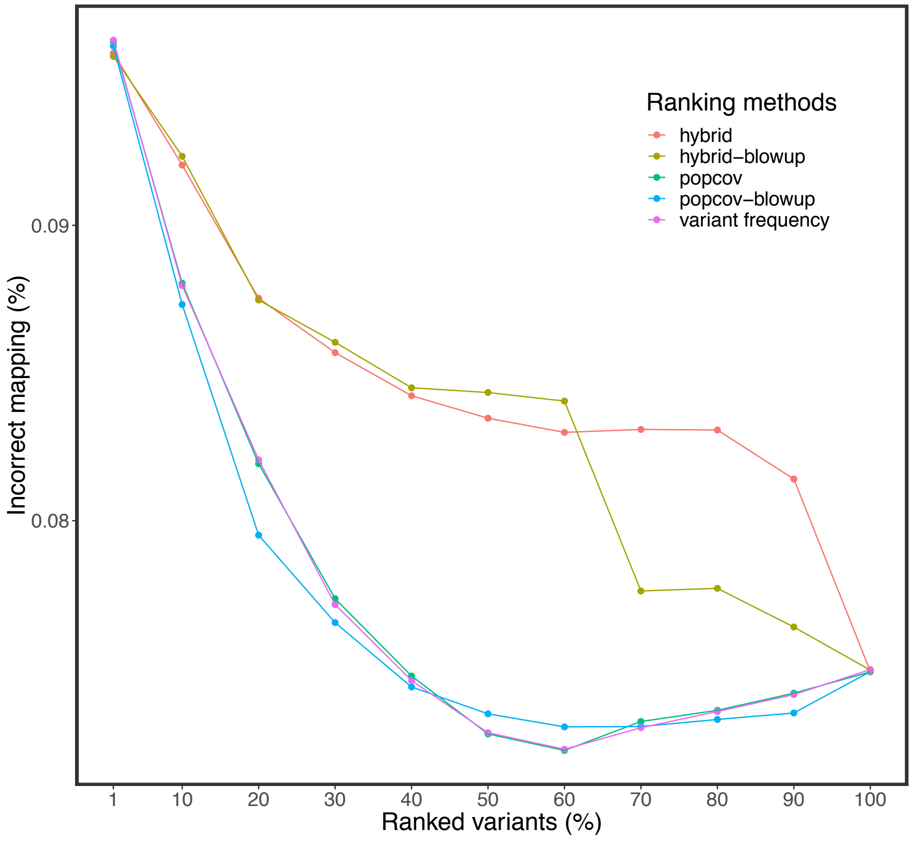


**Figure SN1: Comparison of different variant prioritization strategies.**
Proportion of incorrectly mapped reads for graphs constructed with five variant prioritization approaches.

### Supplementary Note 2

### Adjusted (tuned) linear mapping approach

We followed the proposed approach outlined by Grytten et. al., (2020) to adjust the default parameters of *BWA mem* in order to also consider sub-optimal alignments.
First, we reduce the D value (default 0.5) to consider more alternative alignment positions. However, the mapping performance changed only marginally.
Second, we ran Minimap2 in short read mode (-ax sr) to find all suboptimal alignments. Subsequently, we retained for each read the read placement from either BWA mem or Minimap2 that had the higher alignment score. For reads that had identical alignment score and position for both linear mappers, we retained the lower mapping quality score. For all other cases, we retained the BWA mem alignment.

We made two observations (Figure SN2):

1. The overall mapping accuracy increased mainly due to a smaller number of incorrectly placed reads that had high mapping quality (MQ > 10). This indicates that the tuned linear mapping approach assigns the quality of the alignments better.
2. We found an improvement in mapping accuracy only on reads that are identical to the reference, but not on reads that contain variants.

While Grytten et al. observed that an adjusted parameter setting of BWA mem and subsequent application of Minimap2 led to considerable accuracy improvements, the gain in accuracy was low in our study. The proportion of simulated reads with variants was twice as high (19.16% vs. 10.6%) in our study than in Grytten et al., because the average number of polymorphic sites per genome was almost two-fold higher in cattle than humans (see Additional file 3: Table S1).

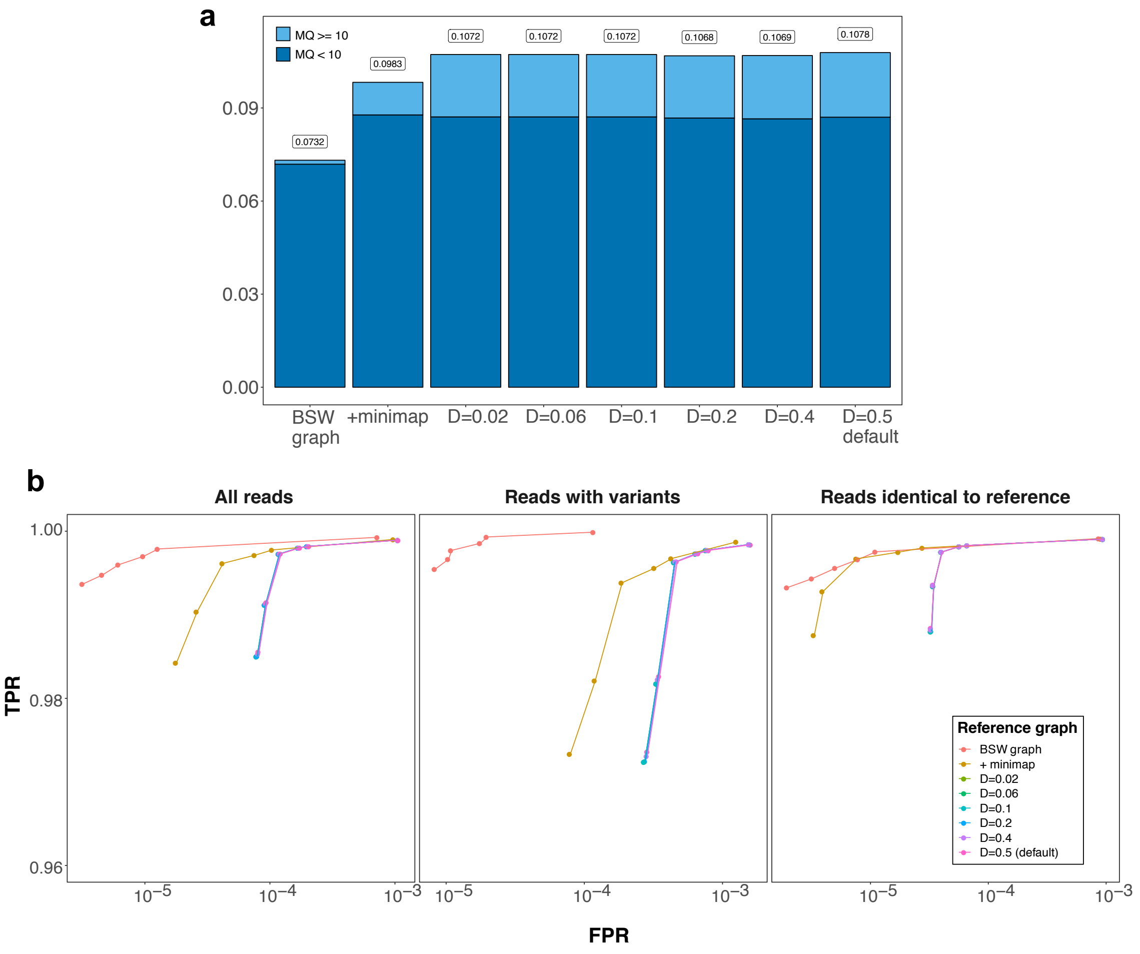


**Figure SN2: Mapping accuracy of paired-end reads simulated form a Brown Swiss animal using different mapping approaches.**

(a) Proportion of simulated reads with mapping errors for different mapping scenarios. (b) True positive and false positive rate parameterized on mapping quality for the different scenarios.

### Supplementary Note 3

### Integrating structural variants into the graphs

We investigated the effect of including longer (structural) variants. For this purpose, we first called and genotyped structural variants using *Delly* (Rausch *et al.* 2012) from 82 Brown Swiss samples that had been sequenced using short-reads (see Material and Methods part of the main manuscript). We discovered 157 precise SVs on bovine chromosome 25 that had an average length of 178 bp. We then combined these variants with 243,145 SNPs and Indels that were discovered using GATK. We used the bta25 ARS-UCD1.2 reference as a backbone and constructed four graphs: (i) SNPs (+Indels) from GATK, (i) SVs from Delly, (iii) SNPs (+Indels) from GATK + SVs from Delly, (iv) empty (only the backbone, no variants). We simulated 10 million paired end reads from haplotypes of one Brown Swiss animal (SAMEA6272105, that had 121,996 SNPs + Indels and 57 SVs that were included in the graph). The simulated reads were mapped to the different graphs using vg.

**Table SN3: Mapping accuracy for graphs that contained different variant types**
MQ=0 and MQ < 10 indicates the proportion of reads mapped with mapping quality 0 and less than 10, respectively.

| Graphs | Variants in the graphs | MQ=0 (%) | MQ<10 (%) | Mapping error (%) |
|---|---|---|---|---|
| Linear | 0 | 0.15474 | 0.22310 | 0.08599 |
| SNP | 243,145 | 0.15366 | 0.21804 | 0.07995 |
| SV | 157 | 0.15508 | 0.22390 | 0.08629 |
| SNP + SV | 243,145 + 157 | 0.15458 | 0.21900 | 0.08003 |

Adding SVs that were detect from short sequencing reads to the graph marginally affected the mapping performance. Actually, the mapping accuracy decreased slightly when SVs were added. Read mapping accuracy improvements were attributable to the SNPs and Indels detected using GATK.
